# Recording neural activity in unrestrained animals with 3D tracking two photon microscopy

**DOI:** 10.1101/213942

**Authors:** Doycho Karagyozov, Mirna Mihovilovic Skanata, Amanda Lesar, Marc Gershow

## Abstract

Optical recordings of neural activity in behaving animals can reveal the neural correlates of decision making, but such recordings are compromised by brain motion that often accompanies behavior. Two-photon point scanning microscopy is especially sensitive to motion artifacts, and to date, two-photon recording of activity has required rigid mechanical coupling between the brain and microscope. To overcome these difficulties, we developed a two-photon tracking microscope with extremely low latency (360 μs) feedback implemented in hardware. We maintained continuous focus on neurons moving with velocities of 3 mm/s and accelerations of 1 m/s^2^ both in-plane and axially, allowing high-bandwidth measurements with modest excitation power. We recorded from motor- and inter-neurons in unrestrained freely behaving fruit fly larvae, correlating neural activity with stimulus presentation and behavioral outputs. Our technique can be extended to stabilize recordings in a variety of moving substrates.

## Introduction

To understand how the brain selects and enacts behavior, it is desirable to record activity in behaving animals. Optical recording of neural activity has become a standard technique in systems neuroscience, but making these measurements in freely behaving animals poses technical challenges[1, 2]. Behavior is expressed as motion, and motion of the brain limits the accuracy of optical imaging techniques. Fundamentally, optical measurement of neural activity requires precisely measuring the amount of light emitted by a fluorescent indicator. Movement of a labeled neuron relative to the microscope objective will alter the efficiency with which the indicator is excited and with which fluorescence emissions are collected, as will changes in the position or properties of scattering elements between the objective and the neuron. If these changes or movements accompany the animal’s behavior, the result will be a fluorescence signal that varies due to motion, not neural activity.

The most common solution to the problem of brain motion is to rigidly couple the brain to the objective. For example, mice[3] and adult flies[4–7] can be head-fixed to the objective while exploring virtual environments controlled by motion of the animal’s legs or wings, or the microscope itself can be mounted on a behaving rodent[8–11]. Larval zebrafish have been paralyzed and embedded in agar with fictive motion read out through electrical recording of the motor neurons[12]. Whole brain imaging has been accomplished without simultaneous measurement of behavior using a variety of techniques in immobilized *C. elegans* [13–15] and with light-sheet microscopy in dissected *D. melanogaster* larval brains[16].

Small transparent genetic model organisms, like the nematode *C. elegans* and the larval stages of the fruit fly *D. melanogaster* and zebrafish *D. rerio* are particularly suited to optical interrogation, because the indicators may be genetically targeted to the neurons of interest and the entire nervous system is optically accessible without surgery or implantation. These organisms offer the possibility of recording from intact, unrestrained, and freely behaving animals, using feedback to keep the brain centered in a larger imaging volume and ratiometric techniques to correct for motion artifacts. This approach has been applied most successfully in *C. elegans*, with wide-field fluorescence or spinning-disk confocal microscopy[13, 17–20].

In other transparent organisms, a modified light sheet technique has been used to locate the positions (but not yet the activities) of neurons in moving *Drosophila* larvae[21]. Spinning disk confocal microscopy was used to detect bilateral differences in activity among populations of neurons in newly hatched crawling *Drosophila* larvae[22]. Recently, extended depth of field microscopy has been used to record from neurons in moving zebrafish[23], and neural activity has been measured using a wide-field fluorescence microscopy in *Hydra* confined to a plane [24]. All these measurements used single photon fluorescence excitation. This can be problematic, because scattering limits imaging in thicker and less transparent models and because the excitation light complicates the simultaneous use of optogenetic reagents while providing an unwanted visual stimulus (except to worms, which have a limited light response[25]).

Here we report a two-photon microscope capable of recording neural activity in freely and three-dimensionally moving intact semi-transparent animals. We applied this microscope to the study of larval *Drosophila*, an attractive model in which to study circuits that mediate decision making [26–32]. The larva has a legion of advantages for systems neuroscience - an optically accessible and representative insect brain, simple behaviors, an emerging wiring diagram[33–36], and powerful genetic reagents that sparsely label neurons throughout the brain[37, 38]. Recording neural activity from behaving larvae has proved elusive, due to scattering in the cuticle and internal viscera and to peristaltic contractions that jerk the brain internally forward and backward and, especially challenging for microscopy, up and down as well[39, 40]. We overcome these challenges, recording from targeted motor- and inter-neurons in freely behaving second instar larvae, including visually responding interneurons and pairs of neighboring neurons. This work presents the first measurements of activity in behaving larvae and the first two photon measurements of activity in any animal not rigidly coupled to a microscope objective.

## Results

### Construction and testing of tracking microscope

Recording from three-dimensionally moving neurons requires at a minimum the ability to rapidly sample an extended volume. We first constructed a point-scanning volumetric two photon microscope(Figure 1a), using an ultrasonic acousto-optic lens (TAG lens) as a 70 kHz resonant axial (z-) scanner[41]. However, the ultimate goal of this volumetric imaging was not to form an image of the neurons but instead to reveal their activities. We therefore decided that instead of imaging an entire volume of moving brain, we would track and record only the cell bodies of selected neurons. Continuous tracking greatly increases both the sampling frequency and magnitude of the fluorescence signal recovered from a targeted neuron, allowing high-bandwidth optical measurements that, in principle, rival the temporal resolution of electrophysiology.

**Figure 1:**
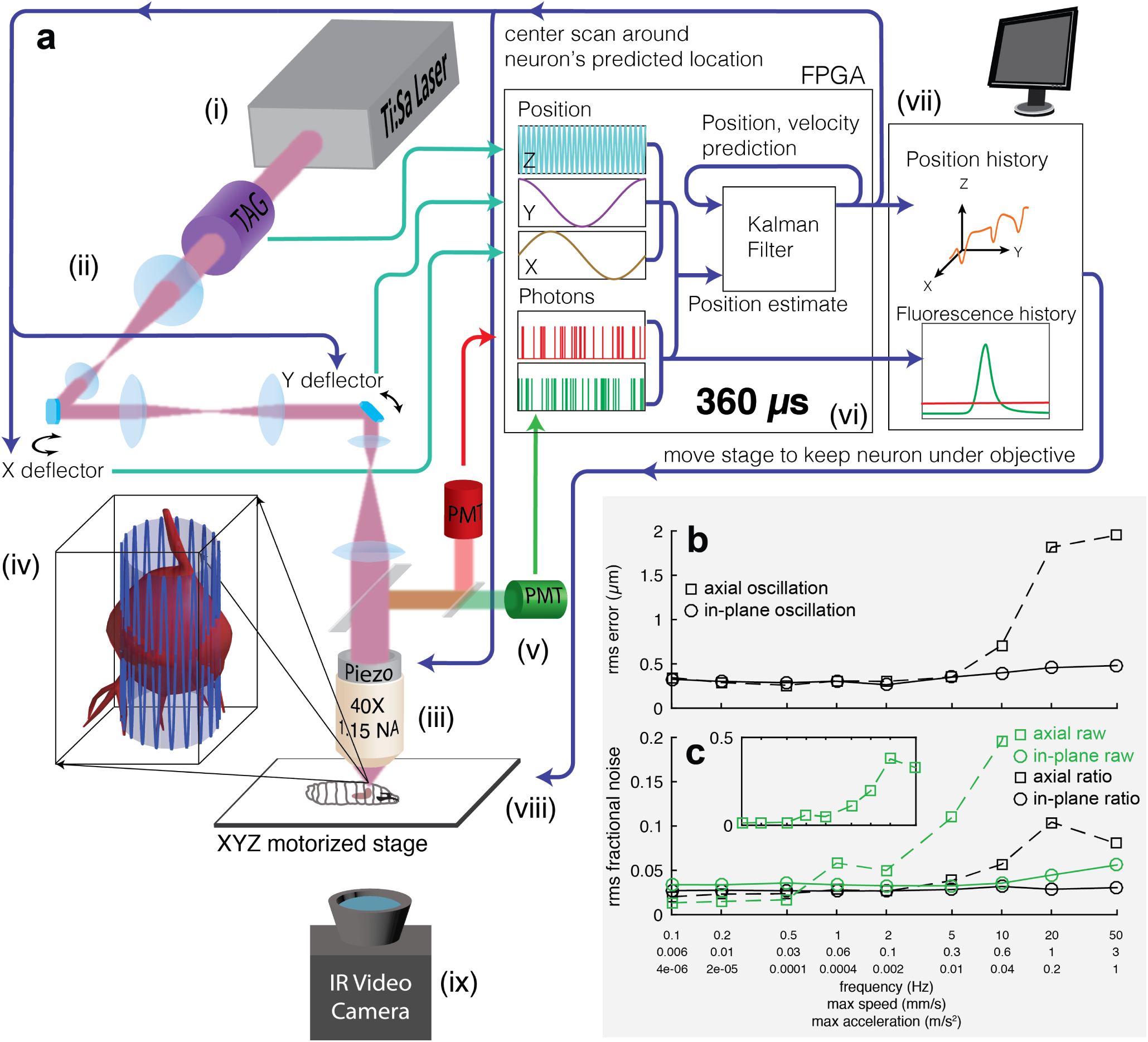
Tracking a moving neuron. (a) Schematic of apparatus. Pulsed infrared excitation light is provided by a Ti:Sa laser(i). Scan optics consisting of two galvanometric mirrors and a resonant ultrasonic lens(ii) shape a wavefront that is relayed on to the back aperture of a piezo-mounted 40x objective(iii), scanning the two-photon excitation focal spot in a cylindrical pattern about a targeted neuron(iv). The scan pattern, schematized by the lightly shaded cylinder and oscillating blue line, has a diameter (7-8 μm) about 75% of that of the targeted neuron. The height of the cylinder (37 μm) has been shortened and the number of z-oscillations reduced for clarity. Red and green fluorescence emission is captured by the objective(iii) and focused through dichroic beamsplitters on to separate photomultiplier tubes(v). Software on a field-programmable-gate-array (FPGA) (vi) correlates photon emission with focal spot position to create an estimate of the neuron’s location. This estimate is combined with previous estimates by a Kalman Filter to produce a prediction of the neuron’s position and velocity. The predicted center location is updated every 360 μs, and each cylindrical scan is centered about the neuron’s newly predicted location. The predicted position of the neuron and the number of red and green photons counted are sent to a computer(vii) which records data to disk for later analysis and controls a 3-axis stage(viii) to return the neuron to the natural center of the imaging system. A low magnification IR video camera placed below the stage (ix) records the posture of the larva for behavioral analysis. **(b,c)** Tracking error and noise in fluorescence vs. speed and acceleration. A single neuron was tracked while being oscillated in a 20 μm peak-to-peak sinusoidal wave at varying frequencies, either in the focal plane or axially. Note logarithmic x-axis.

We began by tracking and recording continuously from the cell body of a single neuron[18]. In larval *Drosophila*, activity in a single neuron can have profound behavioral consequences[30, 32, 42–45], and no method currently exists to reveal this activity in behaving animals. The technique we developed for single neuron tracking could then be extended to record quasi-simultaneously from multiple neurons and to stabilize imaging in moving volumes.

To track a single neuron, we adopted the “tracking FCS” technique from single molecule biophysics[46, 47]. To follow a biomolecule in two dimensions, tracking FCS scans a laser spot in a circle around the putative location of a fluorescently labeled target. The rate of photon emission at each point on the circle is used to calculate an estimate of the target’s location as well as an uncertainty in that estimate. This estimate is combined with the previous ones to form a new best estimate of the target’s location, and the next circle is executed about this updated location. Several approaches can be used to extend this method to 3 dimensions[48, 49]. We used our TAG lens resonant axial scanner in combination with galvanometric mirrors to create a quasi-cylindrical scan pattern (Figure 1a), an approach recently and independently demonstrated by another group working on molecular tracking[50].

Figure 1a shows a schematic of our tracking scheme. We moved the focus of our pulsed excitation laser in a cylinder 7-8 μm in diameter and 37 μm in height through the soma of the cell. The pulsed laser wavelength of 990 nm excited both GCaMP6f[51], a green indicator of neural activity, and hexameric mCherry[52], which provided a stable red baseline for tracking and ratiometric correction. We directed spectrally separated emission from the two proteins to two PMTs operating in photon-counting mode. We correlated the photon emission with the position of the focal spot and used the spatial map of fluorescence emission to calculate both an estimate of the neuron’s position and an uncertainty in that estimate (methods). We used these estimates to update a Kalman filter[53, 54] to form the best current estimate of the neuron’s position and velocity given all previous measurements and their errors. We then centered the next cylindrical scan about the neuron’s predicted center location. Each cycle took 360 μs, and the entire process was implemented in hardware using a field programmable gate array (FPGA). To maximize the reliability of tracking and minimize variations in measured fluorescence, we used a piezo objective scanner to maintain the neuron’s axial position near the natural focal plane of the objective. A final slower feedback loop with a latency of at least 25 ms used a 3-axis stage to return the neuron to the center of the field of view. An infrared camera beneath the stage recorded the larva’s motion for behavioral analysis.

To test the tracking device, we created a larva expressing hexameric mCherry and hexameric GFP[52] in aCC and RP2 motor neurons[55, 56] in the larva’s ventral nerve cord (VNC). We immobilized the larva and used a 3-axis piezo stage to oscillate it in a sinusoidal motion with 10 μm amplitude and varying frequencies while tracking a single neuron. Our goal was to test the efficiency of the tracking algorithm for rapid movements, so we disengaged the slower stage feedback loop.

We first measured the spatial accuracy of the tracker, quantified as the root-mean-squared (rms) error in the center location of the neuron vs. frequency of oscillation for x- (in plane) and z- (axial) sinusoids(Figure 1b). For in plane motion, the rms error ranged from 300 – 500 nm up to 50 Hz, the highest frequency we could achieve with our piezo stage. At this frequency, the neuron’s peak speed was over 3 mm/s and the peak acceleration was 1 m/s^2^, but the tracker was able to follow the neuron’s motion within <10% of the cell-body diameter. For out of plane motion, the tracking performance began to degrade at speeds exceeding 0.3 mm/sec, but even at the highest speeds, the tracker never lost the neuron and remained within the soma.

Next, we asked how much noise tracking the neuron would add to a measurement of neural activity. Signals from the GCAMP6 family of calcium indicators[51] typically create changes in fluorescence on the order of 100%, so we wanted the noise to remain well below this. We first calculated the rms noise in GFP emission in a 0.1-100 Hz bandwidth, divided by the mean emission (Figure 1c). For in-plane motion, the fractional rms noise ranged from 3 to 7%, of which 2% was due to shot noise, a fundamental limit. For axial-motion, the fractional rms noise increased rapidly for oscillations above 5 Hz, correlating with the increasing positional tracking error. We asked whether ratiometric measurement could correct these motion artifacts. We used a stochastic point process smoother[57] (spps, methods) to find the time-varying ratio of green to red fluorescence; with our choices for spps constants, the fractional rms noise in the green/red ratio remained below 5% for in-plane motion and 12% for axial motion. When we examined the power spectra (Figure S1, Figure S2), we found that for all motions, the ratiometric measurement was shot noise limited, except at the frequency of motion and its harmonics.

Tracking multiple neurons (discussed later) requires the ability to rapidly move the focal spot from one neuron to the next. A limit of our galvanometric microscope is that these transitions cannot be achieved quickly, especially for distant neurons. With non-inertial scanners (e.g. acousto-optic deflectors), this constraint is relaxed, as any two spots within the focal volume of the objective are equally accessible. Using these scanners, our ability to track multiple neurons would be limited by how frequently an individual neuron must be sampled to maintain a lock. To determine this limit, we disabled photon counting and feedback for a progressively larger fraction of the time (Figure S3). We found that using only 10% of the available time, we were able to maintain a lock on a neuron moving up to 0.6 mm/s while keeping the ratiometric noise within acceptable bounds for calcium indicators.

### Recording from a motor neuron in a crawling larva without motion artifacts

To record activity in behaving larvae, we needed to temporarily immobilize the larva to locate the neurons we wished to track before releasing the larva to allow free motion while recording from the targeted neuron(s). We created a microfluidic device (Supplemental Movie 1, methods) that used vacuum to reversibly hold the larva[58]. When the vacuum was engaged, the larva was compressed upward against the coverslip, preventing it from moving. When the larva was released to allow motion, residual compression held the larva’s dorsal surface against the coverslip to improve optical access; this interaction slowed the larva’s motion modestly but otherwise allowed normal movements like forward and backward crawling and head sweeps / body bends.

To begin, we recorded from motor neurons in the larva’s VNC (either aCC or RP2; methods). In dissected[16, 55, 59] or immobilized (Supplemental Movie 2) animals, these neurons show patterned waves of activity that could represent fictive peristalsis, but at a much lower frequency than actual peristaltic crawling, presumably because needed proprioceptive feedback is missing. We reasoned that if we recorded from these neurons in moving animals, we should see activity that was timed with peristaltic crawling. Indeed, when we recorded from a motor neuron (Figure S4) labeled with GCaMP6f[51] in a crawling larva(Figure 2a and Supplemental Movie 3), we found clear peaks of activity coincide with each burst of forward motion.

**Figure 2:**
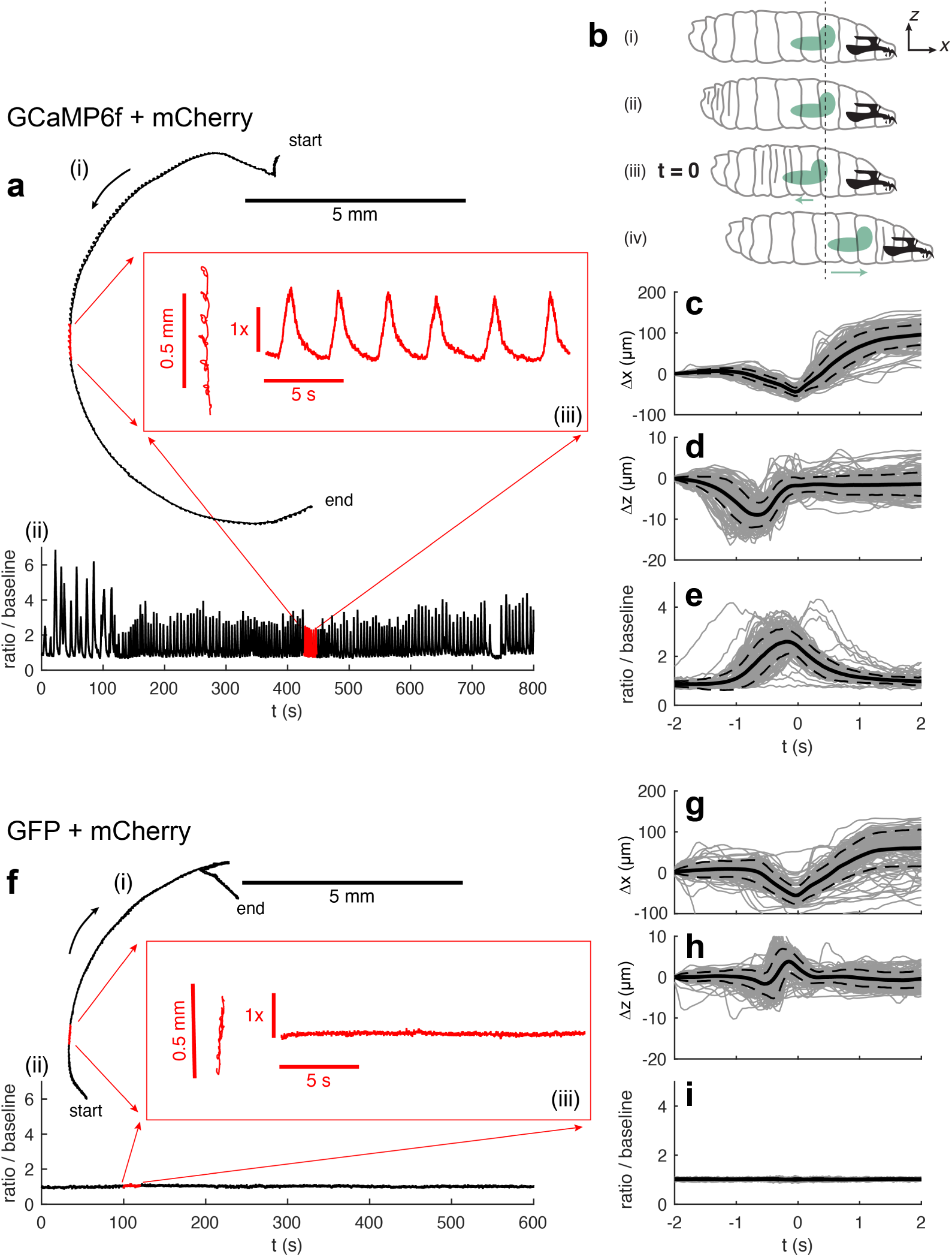
Single motor neuron activity in a crawling larva. **(a)** Path (i) and activity (ii) of a motor neuron labeled with GCaMP6f and hexameric mCherry. For the first 110 seconds, the larva was crawling backwards. Red inset (iii) shows an expanded view of the path and activity during indicated 20 second time interval. **(b)** Sketch of larva’s peristaltic cycle[40]. (i) initial position of the larva & reference coordinate system. (ii) The larva initiates a peristaltic wave with a posterior contraction. (iii) As the wave passes forward, the brain first moves backwards towards the posterior. In **(c-e)** and **(g-i)** the time at which the brain is furthest back is set to the reference time t = 0. (iv) The brain and anterior of the larva move forward together as the larva completes the peristaltic cycle. **(c-e)** Position and ratiometric activity measure of the neuron vs. time in peristaltic cycle. t = 0 represents the point in each cycle at which the brain is furthest back. Individual traces are shown in light gray; the mean trace is in solid black and the dashed lines represent one standard deviation from the mean. n = 135 peristaltic cycles **(f-i)** The same measurements as shown in **(a,c-e)**, but for a motor neuron labeled with hexameric GFP and hexameric mCherry. n = 130 peristaltic cycles.

If the motor neuron’s activity is indeed responsible for coordinating peristalsis, we expect the activity to be repeatable and phase-locked to the motion. With each peristaltic cycle the brain first moves backwards (and down slightly) before accelerating forwards[40]. We chose the point at which the brain is furthest back as a marker of a particular point in the peristaltic cycle(Figure 2b) and aligned both the motion of the neuron and the ratiometric activity measure to this time point (Figure 2c-e). The activity is coherent and phase-locked to the forward motion.

Although our calibration measurements (Figure 1) show minimal motion artifacts when the entire larva is translated as a body, peristaltic crawling moves and deforms the brain as well as the cuticle and the internal viscera between the brain and objective. It is reasonable to wonder if the observed changes in fluorescence ratio were due to calcium transients in the neuron or to an artifact associated with motion. As a control, we therefore made similar measurements in larvae whose motor neurons were labeled with hexameric GFP and hexameric mCherry. GFP fluorescence is not sensitive to calcium concentration, so if the oscillations we observed in GCaMP6f expressing neurons reflected calcium dynamics, we expected these oscillations to be absent in GFP expressing neurons. Indeed, we found the ratio of GFP/mCherry emission to be stable throughout the movement (Figure 2f and Supplemental Movie 4).

We aligned the motion and ratiometric measure of the GFP/mCherry neuron during each peristaltic cycle to the point in time when the brain was furthest back (Figure 2b). We found repeated movements (Figure 2g,h) that were similar but not identical to those in the GCaMP6f expressing larva (Figure 2c,d), an expected result when comparing the fine details of motion between different larvae, but the GFP/mCherry ratio did not vary with this movement (Figure 2i).

### Sequential recordings from motor neurons in the same crawling larva reveal activity timed to behavior

Next we explored how the timing of activity in VNC motor neurons varied with position. We recorded serially from several motor neurons in the same animal. We first immobilized the animal and locked on a neuron. While the larva was immobilized, we used higher power in the excitation laser to photobleach neurons surrounding the targeted neuron, making the neuron clearly identifiable in epifluorescence imaging and preventing the tracker from jumping between adjacent neurons. We then released the immobilization to record both behavior and activity. After recording activity in the neuron for a period of forward movement, we again immobilized the animal, chose a second neuron, and repeated the cycle, for four neurons in total (Figure 3a, Figure S4).

**Figure 3:**
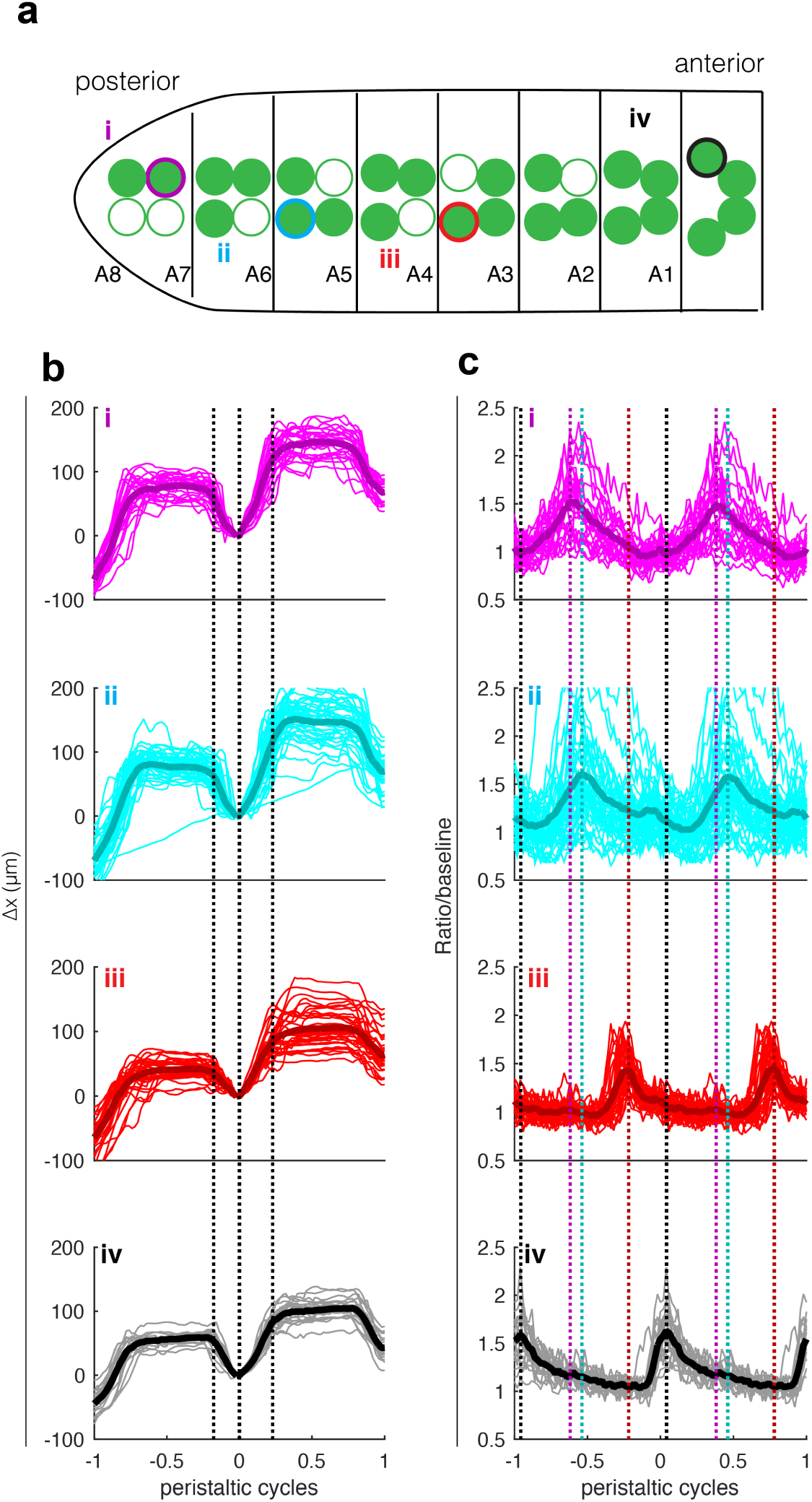
Serial measurements of motor neuron activity throughout VNC of single larva. (a) Sketch showing the locations in the VNC of motor neurons chosen for tracking – color and roman numeral match the labels in (b-c). A8-A1 label segments based on observed positions of cell bodies. Open circles indicate cell bodies were not visible in epifluorescence image. **(b-c)** Position of the neuron along the direction of travel **(b)** and ratiometric activity measure**(c)** aligned to peristaltic cycle. On the horizontal axis, 0 represents the time at which the brain was furthest back in a given cycle, and -1 and +1 represent the times at which the brain was furthest back in the previous cycle and next cycle respectively. Intermediate times were found via spline interpolation. The typical peristaltic period was 6-10 s. Light lines represent individual traces and thick lines represent the mean. In **(b)** black dashed lines show the points of maximal average acceleration for all traces together and are meant to highlight the stereotypy of the aligned motion pattern between traces. In **(c)** the colored dashed lines show the points of maximum activity for the corresponding traces and are meant to aid comparison of the peak locations between traces. From top to bottom, n = 26, 43, 35, 20 peristaltic cycles.

After recording the positions and activities of these neurons, we sought to determine whether they reliably fired at different points in the peristaltic cycle. As for the single neuron experiment of Figure 2, we aligned the temporal axis with t = 0 at the point in the peristaltic cycle where the brain was furthest back. Over the course of repeated immobilizations and releases, the larva varied its peristaltic period, perhaps due to changes in residual compression or motivation to move. We expected both the motion and the activity to be more naturally tied to the local peristaltic period than to a global clock, so to allow comparison between neurons as this period varied, we further aligned the temporal axis so each peristaltic cycle took 1 unit of time. When we performed this alignment, we found each of the 4 neurons moved in the same stereotyped pattern (Figure 3b), but that each neuron was most active at a different point in the peristaltic cycle, with more posterior neurons active earlier (Figure 3c). Finally, we aligned playback of the video recordings of the behavior to the clock generated by the neurons’ motions (Supplemental Movie 5). With this alignment, the peristaltic waves of muscle contraction visible in the video are synchronized and the pattern of activity in the VNC motor neurons can be seen to progress from posterior to anterior.

### Simultaneous recording from two motor neurons reveals exact timing differences and confirms tracker’s spatial accuracy

Of course, to tease out timing differences between neurons, it would be preferable to record from them simultaneously rather than rely on synchronization via a behavioral clock. To follow two neurons at a time, we programmed the microscope to execute four cylindrical tracking cycles around a neuron to update its position and sample its activity, then move to a nearby neuron and follow it for four cycles before jumping back. We allowed two cycles (720 μs) for travel time between the two neurons, so the tracker was active 2/3 of the time, 1/3 on each neuron. We programmed the stage and piezo feedback to return the midpoint between the neurons to the natural focus of the objective. Figure 4a and Supplemental Movie 6 show recordings of position and activity made from two VNC motor neurons (Figure 4b, Figure S4) in a larva crawling forward. To confirm the quality of the tracking, we measured the distance between the two neurons. If the tracker failed to maintain a constant lock on the two neurons and instead jumped to another, off-target, neuron, we would expect the distance between the tracked neurons to suddenly change. If the tracker imprecisely found the positions of the two neurons, we would expect the distance between them to vary by an amount corresponding to the error in location. In fact, we found the distance between the two neurons remained nearly constant (mean 17.0 μm, standard deviation 0.4 μm), indicating the same two neurons were tracked continuously and precisely (Figure 4a). 400 nm rms error in the separation of two neurons is entirely consistent with the 300 nm rms error in the absolute location of a neuron measured in Figure 1b.

**Figure 4:**
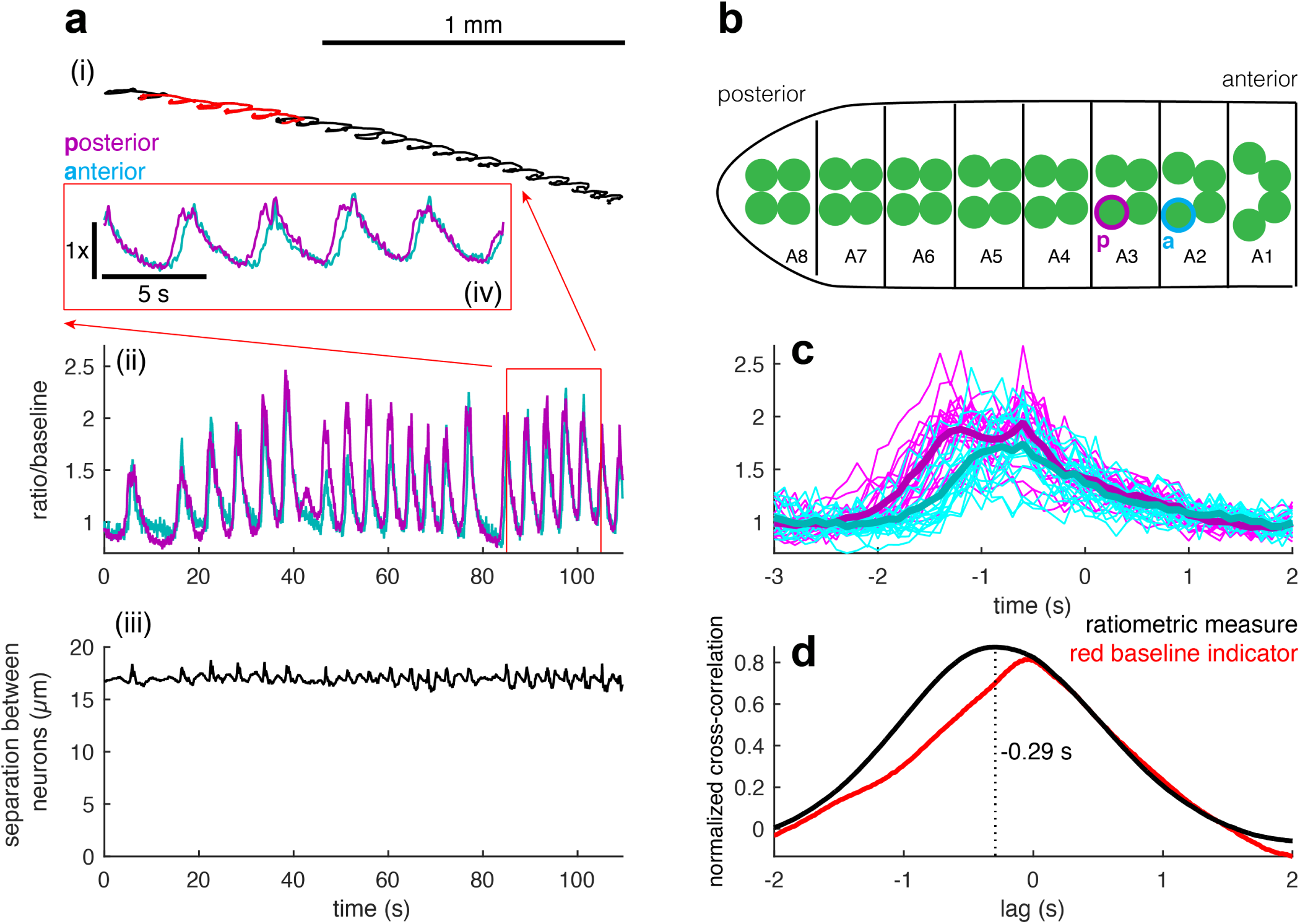
Simultaneous tracking and measurement of activity from motor neurons in adjacent segments. (a) Trajectory of the midpoint between two tracked neurons (i), ratiometric activity measure (ii) for each neuron, and measured distance between neuron centers (iii) vs. time. The inset (iv) shows the activity measures for the two neurons for the highlighted portion of the track. The neuron whose activity is represented by the magenta traces is posterior to the neuron represented by cyan traces as shown in **(b)** a sketch indicating the positions in the VNC of the two tracked neurons. **(c)** Activities of the two neurons temporally aligned to the peristaltic cycle (light lines – individual traces, thick lines – mean). As in Figure 2e, t = 0 when the neuron is furthest back along the direction of travel (n = 20 peristaltic cycles). **(d)** Normalized cross-covariance between the activities (black line) and mCherry emissions (red line) of the posterior and anterior neurons. The ratiometric calcium measure of the posterior neuron leads the anterior by 290 ms, while there is no lag between the mCherry signals.

When we aligned both neurons’ activities to the peristaltic cycle (t = 0 when the brain was furthest back as shown in Figure 2b), we found that the more posterior neuron was active earlier (Figure 4c) in the cycle. To confirm that the posterior neuron led the anterior neuron, we calculated the normalized cross-covariance (methods) between the two neurons’ activities (Figure 4d). The location of the maximum of the cross-covariance tells the delay between the two signals. We found a maximum at *τ* = −0.29 s, indicating activity in the posterior neuron led the anterior by 290 ms. As a control, we calculated the normalized cross-covariance between the red fluorescent signals of each neuron and found the peak correlation to be at zero lag, showing that the ratiometric measure of activity in the posterior neuron leads the anterior neuron because the calcium transients in the posterior neuron lead those in the anterior and not due to some motion artifact.

### Light-evoked activity in an interneuron in the visual pathway

Behavioral responses to stimuli are often variable, even when the stimulus itself is precisely repeated. Understanding how variability in behavior arises from variability in neural activity requires simultaneous measurement of the activity and behavior. Larval *Drosophila*’s variable response to visual stimuli has been studied in the context of uni- and multi-sensory decision making[42, 60–64]. Because the exquisitely sensitive visual system is located in close proximity to its brain[60–62, 65]; optically probing visual circuits in behaving larvae requires multiphoton tracking microscopy.

The “5th LaN,” an interneuron innervating the larval optical neuropil[35, 66] and required for light avoidance responds to blue light presentation to the larval visual organ[60, 62]. To demonstrate our microscope’s ability to record activity encoding visual cues in behaving animals, we recorded from the 5th LaN while presenting short pulses of blue light (methods) to a freely moving larva (Figure 5a, Supplemental Movie 7, Figure S4). We expect that because the 5th LaN is early in the visual pathway, its activity should encode the light stimulus and not motor output. Indeed, when we align the activity of the neuron to its motion (as in Figure 2), we find little variation in the mean activity vs. the time within the peristaltic cycle (Figure 5b), showing that the measured activity of the neuron does not reflect the larva’s peristaltic motion. Aligning the activity of the neuron to the onset of the blue light stimulus, on the other hand, (Figure 5c) shows a consistent response to light presentation.

**Figure 5:**
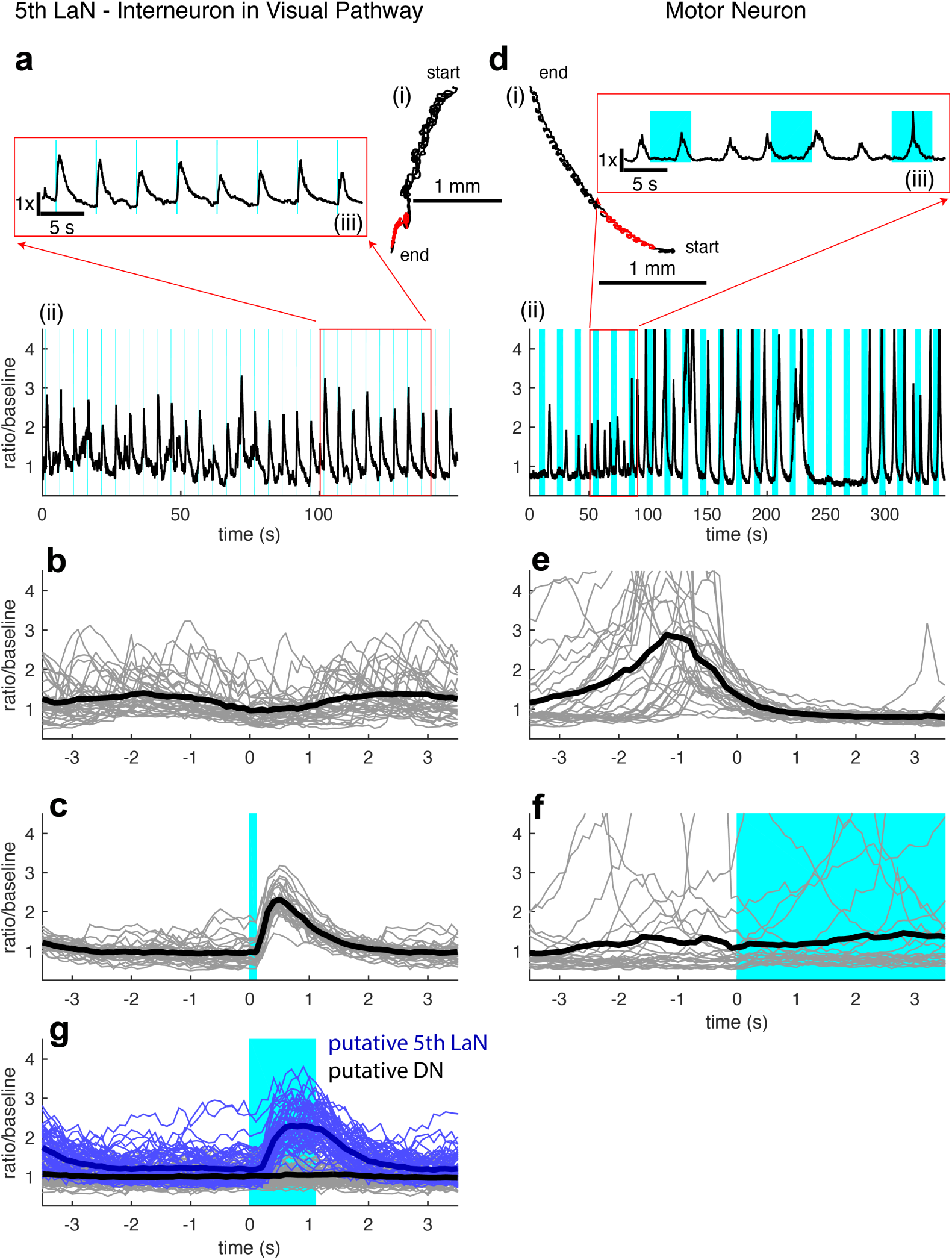
Recording activity from an interneuron in the visual system. (a) Trajectory (i) and ratiometric activity measure (ii) of the 5th LaN. Shaded blue regions indicate times when visual stimulus was presented (100 ms pulses every 5 seconds). The inset (iii) shows the activity for the highlighted portion of the track. **(b)** Activity of the 5th LaN temporally aligned to the peristaltic cycle. As in Figure 2e, t = 0 is the point at which the neuron is furthest back along the direction of travel (n = 35 peristaltic cycles). **(c)** Activity of the 5th LaN temporally aligned to the presentation of blue light stimulus; t= 0 represents the onset of stimulus. Blue shaded region indicates stimulus was on (n = 30 light presentations). **(d)** Trajectory (i) and ratiometric activity measure (ii) of an aCC/RP2 motor neuron. Shaded blue regions indicate times when visual stimulus was presented (5 second pulses every 15 seconds). The inset (iii) shows the activity for the highlighted portion of the track. **(e)** Activity of the motor neuron temporally aligned to the peristaltic cycle. As in Figure 2e, t = 0 when the neuron is furthest back along the direction of travel (n = 30 peristaltic cycles). **(f)** Activity of the motor neuron temporally aligned to the presentation of blue light stimulus; t = 0 represents the onset of stimulus. Blue shaded region indicates stimulus was on (n = 23 light presentations). **(g)** Simultaneous recording in a moving larva from two TIM expressing neurons. As in **(c,f)** recordings are aligned to the onset of blue light stimulus and shaded region indicates time when stimulus was on (n = 61 light presentations). **(b,c,e-g)** Light lines represent individual traces; thick lines represent mean.

The measured response of the 5th LaN to blue light is temporally displaced from the actual blue light presentation and of significantly longer duration (Figure 5c), so it is unlikely that this measurement reflects cross-talk from the stimulus presentation. To confirm that there was no cross-talk, we recorded from a motor neuron in a moving animal, while presenting long (5s duration) blue light pulses (Figure 5d, Supplemental Movie 8). We found that, in contrast to the visual interneuron, the motor neuron’s activity was correlated with motion (Figure 5e) but did not reflect the stimulus presentation (Figure 5f).

Our ability to simultaneously record from neighboring neurons allowed an additional control for cross-talk between the light stimulus and measured activity. We recorded simultaneously from the nearby cell bodies of two neurons labeled by tim-Gal4, one light-responsive and one not. Based on the positions of these two neurons’ cell bodies (Figure S4), we believed them to be the 5th LaN and one of the DN2 neurons, which are known not to respond to light, but the validity of the control does not depend on the specific identities of these neurons. One of the neurons responds reliably to a one second blue light pulse every 5 seconds, while the other shows no activity in response to light stimulation (Figure 5g). Because these recordings were made simultaneously in the same moving animal in response to the same light stimulus, the only explanation for the different behavior of the ratiometric measure is that this measure represents a light-evoked calcium transient in only one of the neurons.

### Activity in a premotor interneuron correlates with behavioral state

We sought to further explore the utility of combined measurements of activity and behavior in the larva. Recent work[59] has identified A27h, an excitatory premotor interneuron present in every hemisegment, as an important component of the circuit coordinating peristaltic motion and also identified a driver line that labels these neurons. Imaging in a dissected preparation showed that activity in the A27h neuron coordinates with waves of motor neuron activity propagating from posterior to anterior but that the neuron is silent when waves propagate in the reverse direction, suggesting these neurons are involved only in forward peristalsis. However, because the imaging was done in an isolated CNS, there remains at least a formal possibility that a different relation might be observed between A27h and the direction of locomotion in an intact crawling larva.

We therefore recorded from A27h interneurons in larvae exploring our microfluidic device. We used sudden changes in the compression applied by the device as a stimulus to encourage transitions between forward and backward crawling and were able to observe multiple transitions while recording from a posterior (A8) A27h interneuron (Figure 6a, Supplemental Movie 9,9, Figure S4). We found that the neuron was active in sync with the peristaltic cycle during forward but not backward crawling, in agreement with the isolated CNS recordings[59]. We then recorded from an A27h interneuron in a more anterior segment (A1) and found that it too was active selectively during forward peristalsis (Figure 6b, Supplemental Movie 11,11, Figure S4). We aligned the activities of the neurons to brain motion during forward peristalsis (Figure 6a,b) and found that the posterior neuron was active earlier in the cycle than the anterior one. Thus, as in the dissected prep, A27h neurons in moving animals encode both the direction of peristalsis and the phase of the peristaltic cycle, supporting the identification of waves of VNC activity in dissected animals as fictive peristalsis.

**Figure 6:**
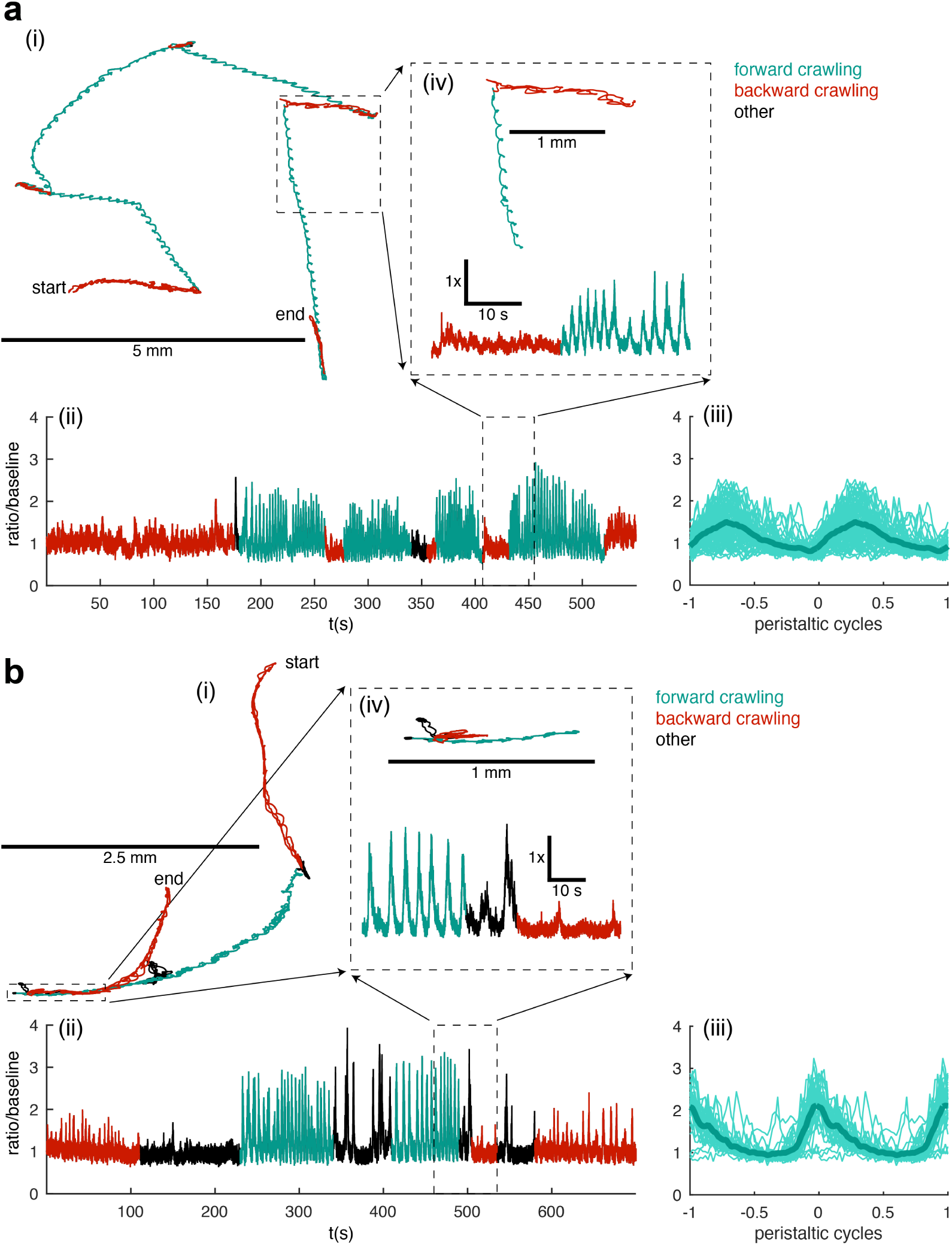
A27h premotor interneurons recorded in larvae crawling forward and backward. (a) Path (i), activity (ii), and activity aligned to forward peristaltic cycles only (iii) of an A27h premotor interneuron located at the posterior of the VNC (A8 segment). Behavioral state of the animal, as identified by inspection of the video recording, is indicated by color: red for backward crawling, teal for forward crawling, and black for other behaviors, like pausing and bending the body without either forward or backward movement. The inset (iv) shows an expanded view of the path and activity for a backward to forward crawling transition (inset time = 48 s, 24 s each backward and forward crawling). n = 80 forward peristaltic cycles **(b)** Path (i), activity (ii), and activity aligned to forward peristaltic cycles only (iii) of a more anterior A27h premotor interneuron (A1 segment). Behavioral state of the animal, as identified by inspection of the video recording, is indicated by color as in **(a)**. The inset (iv) shows an expanded view of the path and activity for a forward to backward crawling transition (inset total time = 75 s, 30 s forwards, 15 s body bend and hunch, 30 s backwards). N = 35 forward peristaltic cycles. **(a,b iii)** Light lines represent individual traces; thick lines represent mean.

## Discussion

Optical recordings in compact and transparent model organisms offer the potential to understand how neural dynamics encode and process information. However, away from the sensory periphery, as neurons’ roles shift from processing sensory input to controlling behavioral responses, activity recorded in an immobilized animal becomes progressively harder to interpret. Here we show for the first time that it is possible to make two photon recordings of neural activity at cellular resolution in an intact freely behaving transparent model organism.

Experiments in the nematode *C. elegans* show the potential to use optical tools to read out patterns of activity in unrestrained freely behaving organisms[13, 17–20], but unique properties of *C. elegans* – high transparency, smooth planar sidewinding locomotion, slow calcium transients in the neurons, and lack of a major visual system – conspire to allow conventional techniques to be used for these measurements. Our novel two-photon tracking microscope will allow measurement in a wide range of behaving visually responsive semi-transparent animals, including larval *Drosophila*, larval *Platynereis*[67, 68] and *Hydra*[24].

In this work, we demonstrated two-photon recording from freely behaving *Drosophila* larvae, a long-held goal in the study of this model organism. The ongoing EM-reconstruction of the larva’s nervous system[33–36], a toolkit of lines that allow sparse labeling of neurons throughout the brain[37, 38], and high resolution behavioral screens have allowed for the discovery of circuit elements well downstream of the sensory periphery[26–32], and it has long been desired to record the activities of these neurons in a behavioral context. However, despite the superficial similarities of *C. elegans* and *Drosophila*, methods that succeeded in the worm have never successfully been used to measure activity in moving *Drosophila* neurons.

For the first time, our microscope allows recording in behaving larvae, enabling significant advances in combination with the larva’s other advantages as a model system. For instance, dopaminergic neurons providing input to the adult mushroom body encode motion among other signals[69], a discovery made by correlating activity and flailing leg movement in a head-fixed fly. Our microscope will allow recording from the corresponding neurons in truly freely moving larvae, and then with the advantage of the EM-wiring diagram and libraries of driver lines, their synaptic partners as well.

The ability to precisely locate fiducial markers in moving tissue has applications beyond measurement of neural activity. In both calibration experiments (Figure 1b) and behaving animals (Figure 4a), we tracked moving neurons with a measured spatial precision of better than half a micron.

### Advantages of two-photon tracking microscopy

To date, our recordings represent the only measurements of neural activity in behaving larvae, but rapid progress in single photon light field and light sheet microscopy gives hope that these methods may soon also be capable of similar measurements. While these approaches potentially allow many neurons to be sampled simultaneously, our two photon tracking approach offers a number of advantages.

#### Two photon microscopy has better penetration and greater compatibility with visual and optogenetic manipulations

As they develop, many transparent animals, including larvae of both *Drosophila* and *Platynereis* [70], become larger and less transparent, rendering two-photon excitation a necessity for imaging throughout the interior. Most of the reagents for sparsely labeling neurons in the larval CNS have been characterized[38] in later developmental stages, so it is highly desirable to conduct experiments in the more developed larvae. Two-photon excitation also helps avoid cross-talk with optogenetic reagents and potential effects of phototoxicity.

Other than worms, most transparent animals have responsive visual systems, and it is impossible to use single-photon excitation to image their brains without also shining visible light in their eyes. As an example, a recent measurement of light-evoked activity in *Hydra* used the onset of fluorescence excitation itself as the sole visual stimulus[24]. Our two-photon tracking microscope will allow the study of neural activity underlying underlying sensory-guided behaviors, including visual and multi-modal navigation, without visible excitation light providing a confounding stimulus.

#### Tracking microscopy increases bandwidth and signal strength

As calcium indicators improve in temporal resolution and genetically encoded voltage indicators enter wider use, new microscopy techniques will be needed to take advantage of the higher bandwidths offered by these indicators. The bandwidth of an optical measurement is limited both by the Nyquist limit imposed by the sampling frequency and by the shot noise limit, which is a function of the raw emission rate of the indicator, the sampling time, and the desired fractional resolution.

Two-photon point scanning microscopy uses detectors with very high bandwidth (>10 MHz for a PMT compared to *<∼*100 Hz for a scientific camera), but traditional two-photon microscopy suffers from low sampling rates and short sampling durations, a result of sequentially sampling millions of individual voxels in a volume that may contain only a few points of interest. This insight has driven the recent development of random access two photon microscopes, which can achieve high bandwidth by recording from only the structures of interest[71–75].

Our tracking approach continuously samples from targeted neuron(s), allowing monitoring with a bandwidth exceeding 1 kHz (the sampling rate is 2.8 kHz for a single neuron, limited by the resonant frequency of the galvo mirrors), a speed compatible with the fastest available voltage indicators. In comparison, SCAPE, a light sheet microscope with improved geometry for measuring behavior, has a volume rate of 10-20 Hz[21], creating a Nyquist limit of 5-10 Hz, below the bandwidth of even existing calcium indicators. Somewhat surprisingly, despite continuously focusing on a single neuron, we did not find photobleaching to interfere with tracking or measuring activity (methods).

### Tracking many neurons and volumetric imaging in moving animals

In this work, we used two photon tracking microscopy to record from up to two neurons in an intact freely behaving transparent animal. Our ability to simultaneously track more than two neurons was limited by the need to physically move the mirrors to move the focal spot from one neuron to the next. However, we showed that we could maintain focus on a single neuron using only 1/10 of the microscope’s tracking time (Figure S3). Using non-inertial scanners, we could therefore expect to be able to track at least 10 separated neurons simultaneously, without assuming any correlations between their motions.

In fact, neurons within a brain have highly correlated motions. If each neuron’s location is considered to be a point in a volume subject to affine transformations (translation, rotation, scaling, and shear), then sampling 4 neurons from a group is sufficient to update the positions of the entire set. A more advanced algorithm would therefore update the estimate of each neuron’s position based on the measurements of every other one as well, and we expect that if such an algorithm were implemented on a random access microscope, it would allow us to follow a large number of labeled neurons as long as they were all contained within the same focal volume and well-separated from each other.

To record from densely labeled tissue or from amorphous neuropil, we would track fiducial markers to establish a stable coordinate system and carry out volumetric imaging with respect to this coordinate system. Tracking many neurons or interleaving tracking and imaging will require random access point scanning. True 3D random access microscopy[71–75] is technically involved, but a hybrid solution using acousto-optic x-y scanning and a TAG lens for resonant z-scanning would be more straightforward to implement.

Our scheme to correct 3D motion during two-photon recording can be applied to other preparations, like head-fixed mice[76, 77], that suffer from motion artifacts. Recent advances in random access two photon microscopy have allowed for rapid sampling of neural activity throughout a focal volume[74, 75]. By selecting only small volumes containing the neurons of interest out of the much larger volume, the rate at which neural activity is recorded as well as the number of photons collected from each neuron is greatly increased. However, random access sampling greatly increases the sensitivity to sample motion. Uncorrected, even micron-scale motion would result in completely missing the intended targets[75]. It would be straightforward to adapt our tracking algorithm to provide real-time three-dimensional motion correction in random access microscopy.

## Conclusion

We demonstrated a two-photon microscope capable of recording from three dimensionally moving motor- and inter-neurons in intact and freely behaving animals. We measured the relation between neural activity and the behavior it controls and the responses of interneurons to stimulus presentation in behaving *Drosophila* larvae. We demonstrated simultaneous tracking of adjacent neurons and argued that, in combination with non-inertial deflectors, our technique can be extended to simultaneous recording from many neurons and from volumes of neuropil. To our knowledge, this demonstrates the first two-photon measurement of neural activity without rigidly fixing the brain to the microscope.

## Acknowledgements

We thank Shy Shoham, Meng Cui, Margarita Kaplow, Marco Tedaldi, and Justin Blau for discussions, and Stefan Pulver and Simon Sprecher for strains. MMS participated in the CSHL Neurobiology of *Drosophila* course with support from a Helmsley Fellowship. This work was supported by NSF Award 1455015, NIH Award 1DP2EB022359, and a Sloan Research Fellowship to MHG.

This article was formatted using a modified Wenneker Article template from LaTeXTemplates.com. This article has not been peer-reviewed or professionally edited. Please contact the corresponding author with any suggestions or corrections.

## Author Contributions

Design, construction, and programming of microscope: MMS, DK, MG. Design of microfluidic devices: MMS, DK, AL, MG. Construction of microfluidic devices: AL. Carried out experiments: MMS, DK. Analyzed data: MMS, DK, MG. Drafted manuscript: MMS, DK, MG. Supervised project: MG

## Methods

### Fly strains

The following fly strains were used:

- w[1118];20XUAS-IVS-GCaMP6f (Bloomington Stock #42747)
- y[1]w[*];Sp/CyO;20XUAS-6XmCherry-HA (Bloomington stock #52268)
- y[1] w[*]; 20XUAS-6XGFP/CyO (Bloomington stock #52261)
- w;+;RRAF-Gal4, UAS-GCaMP6f (gift from Stefan Pulver, University of St. Andrews)
- y[1] w[*];; eve-GAL4.RRK/TM3, Sb (Bloomington stock #42739)
- Cry-Gal80;A3(tim-Gal4) (gift from Simon Sprecher, University of Fribourg)
- GMR36G02-GAL4 (Bloomington stock #49939)
- w[1118]/Dp(1;Y)y[+]; CyO/Bl; TM2/TM6B, Tb (Bloomington stock #3704)

### Crosses

- Figure 1, Figure 2f-i: w; 20XUAS-6XGFP; 20XUAS-6XmCherry-HA (created from Bloomington stock #52261 and #52268 using #3704 balancer) crossed to eve-GAL4.RRK
- Figure 2a-e, Figure 5d-f: w;+;RRAF-Gal4,UASGCaMP6f crossed to UAS-6XmCherry-HA
- Figure 3, Figure 4, Supplemental Movie 2: w;+;RRAF-Gal4,UAS-GCaMP6f crossed to w; 20XUAS-GCaMP6f; 20XUAS-6XmCherry-HA (created from Bloomington stock #42747 and #52268 using #3704 balancer)
- Figure 5 a-c,g: w; 20XUAS-GCaMP6f; 20XUAS-6XmCherry-HA crossed to Cry-Gal80;tim-Gal4*
- Figure 6: w; 20XUAS-GCaMP6f; 20XUAS-6XmCherry-HA crossed to GMR36G02-GAL4 (Bloomington stock #49939)

F1 progeny of both sexes were used for experiments.

*in our lab conditions, we observed gal4 driven expression of transgenes in the Pdf expressing LaNs as well as the 5th LaN due to incomplete suppression by Cry-Gal80.

### Larvae

Flies were placed in vials and allowed to lay eggs for 24 hours at 25°C on standard cornmeal-based food. Second instar larvae, 48-72 hr AEL, were separated from the food with 30% sucrose solution and washed in water. Larvae were hand selected for size and proper expression of fluorescent proteins prior to use in experiments.

### Identification of motor neurons

The lines we used, RRAF-Gal4 and RRK-Gal4 are reported to label aCC and RP2 motor neurons in some conditions and only aCC motor neurons in others[55, 58, 59, 78–82]. We observed labeling of both aCC and RP2 neurons, with some stochasticity in expression. We imaged immobilized RRAF>GCaMP6f,mCherry larvae (same genotype as used in Figure 3,Figure 4) under a fluorescence dissecting scope (Nikon SMZ18, 1.6x objective) and found that all of these neurons were active during fictive peristalsis and during constrained movement (Supplemental Movie 2). We did not attempt to differentiate between aCC and RP2 neurons. We identified the segmental locations of motor neurons based on the position of their cell bodies, assuming the most posterior labeled neurons were in A8 (Figure S4).

### Microfluidic device

In order to allow rapid prototyping and more complex device profiles, we used SLA 3D printing to create microfluidic masters for casting[83, 84]. Masters were designed in Autodesk Inventor and printed on an Ember 3D printer (Autodesk, USA) using black prototyping resin (Colorado Photopolymer Solutions). After printing, masters were washed in isopropyl alcohol, air dried, and baked at 65C for 45 minutes to remove volatile additives and non-crosslinked resin. Baked masters were oxygen plasma cleaned for 10 minutes followed by silanization in a vacuum dessicator with 20 μL of trichloro(1H,1H,2H,2H-perfluoro-octyl)silane (Sigma Aldrich) for at least 4 hours. PDMS (Sylgard 184 dow corning, 10:1 base:cure agent) was poured over the master and baked at 75C for 2 hours. This process resulted in reliable curing and release of PDMS from the master.

The microfluidic device uses vacuum compression to achieve reversible immobilization [58]. The device was designed with three sections: a circular section (diameter 9 mm, depth 200 μm), surrounded by an inner ring (inner diameter 9mm, outer diameter 11.5 mm, depth 100 μm) and an outer ring (inner diameter 11.5mm, outer diameter 13.5 mm, depth 200 μm). The outer ring has a small rectangular channel which is connected via tubing to a vacuum pump. These dimensions were chosen to allow second instar larvae to freely move in the center chamber. The bottom of the PDMS device was bonded to a glass slide for support; a hole was drilled in the slide to allow access for the vacuum connection. A larva was placed into the device along with water for lubrication, then a coverslip was placed on top and held in place using elastic bands (Scunci Girl No Damage Polyband Elastics, Conair Corporation, Stamford, CT). When vacuum is applied to the outer ring, the central chamber is pressed up against the coverslip until the inner ring contacts the coverslip, controlling the compression. When the vacuum is released the larva is free to move, with residual compression serving to keep the dorsal surface in contact with the coverslip (the compression can be fine-tuned by partial release of the vacuum). This process can be repeated without harm to the larva (Supplemental Movie 1).

### Volumetric Two-Photon Microscope

Our volumetric microscope was a custom-built upright microscope with galvanometric mirror-based in plane scanning and resonant axial scanning[41]. Excitation was provided by a tunable Ti:Sa laser with an 80 MHz repetition rate (Chameleon Ultra II, Coherent, Santa Clara, CA); for all experiments described, the excitation wavelength was 990 nm. Beam power was controlled with a Pockels cell (Model 350, Conoptics, Danbury, CT) between crossed polarizers. The power was adjusted for each sample, but the typical power, as measured at the back of the objective was 16 mW. The scan optics consisted of a pair of galvanometric mirrors (8310K, Cambridge Technology, Bedford, MA) separated by a 4f relay (2 lenses: f = 100mm, AC508-100-B-ML, Thorlabs, Newton, NJ) followed by scan (f = 40 mm, AC254-040-B-ML, Thorlabs) and tube (f = 200 mm, AC508-200-ML, Thorlabs) lenses. For all experiments described, a 40X, 1.15 NA water immersion objective was used (N40XLWD, Nikon), mounted on a 100 micron piezo scanner (Nano-F 100S, Mad City Labs, Madison, WI).

To add a resonant z-scan, we placed a TAG resonant ultrasonic lens (TL25*β*.B.NIR controlled by TAG Drv Kit 3.2, TAG Optics, Princeton, NJ) before the microscope in a conjugate plane to the scan mirrors. To determine the exact phase of the lens[41], we sent a 660 nm laser beam (LP660-SF60, Thorlabs) separated from the excitation laser by dicroic beamsplitters (ZT543rdc, Chroma Technology) in the reverse direction through the TAG lens then through an iris onto a photodiode (DET10A, Thorlabs). The power measured by the photodiode varied with the phase of the lens. We fed the photodiode output into a PLL based on the 74HCT9046A (NXP Semiconductors), the output of which we used to determine the phase of the lens and hence the position of the focal spot in z. The phase shift was fine-tuned by matching the images produced on the “fly-forward” and “fly-backward” portions of the resonant cycle. The lens was operated at a frequency of 70 kHz and typically 30% of the maximum amplitude which resulted in a quasi-linear axial scan range (see discussion in volumetric two-photon microscope software below) of approximately 30 μm.

Emitted photons were separated spectrally from the excitation beam by a dichroic beamsplitter, then separated into a red and a green channel by a filter cube containing a dichroic beam splitter and bandpass filters (ZT543rdc, ET510/80m, Chroma Technology and FF02-650, Semrock) and then detected by separate PMTs (R9880U-210 and R9880U-20, Hamamatsu) operating in photon counting mode. These elements were all mounted in the Scientifica Multiphoton Detection Unit (MDU, Scientifica, Sussex, UK). PMT pulses were shaped and converted to digital logic levels by the Hamamatsu Photon Counting Unit C9744. Samples were mounted on a 3-axis motor driven stage (FTP-2000 ASI instruments). The microscope was controlled by a Windows PC and a National Instruments PCIe-7842R multifunction DAQ with on-board FPGA running custom software based on HelioScan (see Volumetric Two-Photon Microscope Software below).

### Epifluorescence imaging

To initially locate and identify neurons prior to tracking, we used wide-field epifluorescence imaging through the same objective as used for two-photon imaging. We removed the long-pass dichroic filter in the MDU from the beam path and inserted a mirror between the scan and tube lens to redirect the light path to travel through a filter cube (either MDF-GFP or MDF-TOM, Thorlabs) and on to a CMOS camera (acA2040-90um, Basler). Depending the fluorophore being imaged, excitation was provided via the filter cube by the collimated output of either a blue (M470L3, Thorlabs) or green (M565L3, Thorlabs) led.

### Bleach mode

To allow unambiguous identification of tracked neurons in epifluorescence images and to avoid the tracker jumping between adjacent neurons in densely labeled tissue, we found it helpful to photobleach the fluorescent proteins in cells near the one(s) being tracked. To do this easily, we added a “bleach mode” to the microscope in volumetric imaging mode. We first selected the neuron(s) to be tracked and then engaged “bleach mode,” which modulated the Pockels cell voltage to image the surrounding volume using a high excitation power (180 mW, as measured at the back aperture of the objective, vs. 16 mW in normal imaging mode) while turning the excitation off completely over the target neurons. Bleach mode was used in advance of tracking for the experiments described in Figure 3 and Figure 4.

### Volumetric Two-Photon Microscope Software

We based our microscope software on the open source Helioscan package written in LabView[85]. As the TAG lens acts as a resonant axial scanner, we rewrote the FPGA-based resonant scanning package to use the z-axis as the fast axis (140 kHz line rate), with the x-axis as an intermediate speed (*∼*1 kHz line rate), and the y-axis as the slowest axis. In order to achieve high volumetric imaging rate, we used all three axes bidirectionally (recording during both the trace and re-trace portion). To overcome the effects of hysteresis, we used the analog outputs of the galvanometer control boards to determine the true position of the focal spot while scanning.

The focus of the system of objective and TAG lens oscillated sinusoidally in time with a frequency of 70 kHz. We divided each oscillation into two scans, one with the focal spot moving away from the objective and one moving towards it. As is standard in resonance scanning, each axial scan was divided into a discrete number of voxels of equal temporal duration, resulting in some distortion due to the nonlinearity of the z-position vs. time. We used the central 60% of the sinusoidal z-scan, meaning that voxels at the extreme of the scan had an axial extent 59% of the voxels in the scan center. For each axial line, the FPGA recorded the x and y position of the galvo (ignoring movement of the galvos during the *∼*4.2 μs z-line duration) and streamed this position along with the photon counts per voxel to the computer. This data was used to assemble image volumes for display or recording. Separate volumes could be formed for the two axial scan directions, allowing us to precisely calibrate a phase delay relative to the TAG lens cycle to bring the two scan directions into alignment.

### Tracking Microscope

The tracking algorithm was based on the tracking FCS method[46, 86]. The FPGA directed the x- and y-galvos to move in a circle of defined radius, centered about the predicted location of the center of the neuron being tracked (taking into account the estimated velocity of motion). The galvos followed this circle faithfully, with a phase-lag that depended on the driving frequency. To correct for the phase-lag, the actual position of the focal spot, as reported by the galvo control boards, was used for all calculations. With each circular scan the FPGA calculated the estimated displacement of the true center of the neuron from the center of the scan (see feedback relations below). To avoid including other cells or autofluorescent background material in the estimate of the neuron’s location or its intensity, only the portion of the resonant scan within *± ∼*8 μm of the estimated center was used for fluorescence measurement.

The FPGA based tracking algorithm reported the location of the neuron within the focal volume and its velocity and used the mirrors and objective piezo to center the cylindrical scan on the neuron. Feedback to the mirrors was updated with every scan, creating a latency of 360 μs. Fast axial re-centering was accomplished by selecting a portion of the TAG axial scan centered on the neuron’s predicted location, again with a latency of 360 μs. Signals to the objective piezo were lowpassed on the FPGA to 70Hz to fit within the bandwidth of the scanner. A separate PID feedback loop running on the computer moved the stage to return the neuron to the natural focus of the objective. Re-centering commands were sent to the stage every 25 ms.

### Feedback Relations

In this discussion and following, *x* and *y* represent displacements in the focal plane of the objective and *z* represents axial displacement. Our cylindrical scan pattern (radius = *r*_*scan*_) was contained radially within the soma and extended axially well beyond the bounds of the neuron. With each cycle, we calculated the sum of all locations where photons were detected and also kept track of the total number of photons.

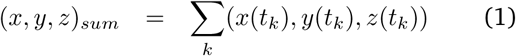

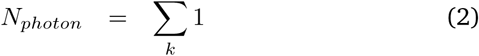

where *t*_*k*_ represents times when photons were detected. We used these sums to estimate (Δ*x,* Δ*y,* Δ*z*), how much the neuron was displaced from the center of the scan, as well as the uncertainty, expressed as a variance, (*R*_Δ*x*_, *R*_Δ*y*_, *R*_Δ*z*_) in this estimate. Derivation of these results, following Berglund and Mabuchi[86], is found in the supplementary methods and assumes that the intensity distribution of the neuron is cylindrically symmetric.

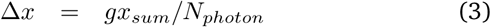

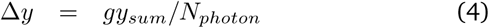

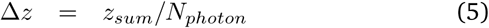

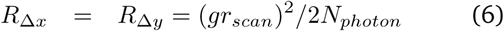

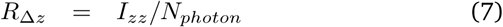

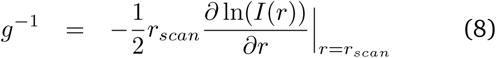

Where *I*(*r*) represents the fluorescent labeling intensity averaged along z at a distance r from the center of the neuron, and 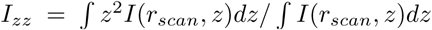 is the second moment of the intensity distribution along the z-axis at the scan radius. *r*_*scan*_ can be chosen to minimize the x- and y-measurement error. Noting that *N*_*photon*_ ∝ *I*(*r*),

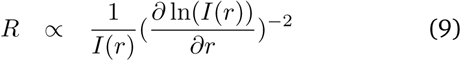

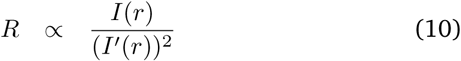

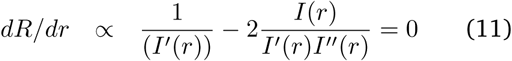

at the minimum error location. For a Gaussian intensity distribution, 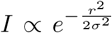, this minimum is found at *r* = 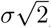, and *g* = 1, recapitulating the result in [86]. If we consider the neuron to be a uniformly labeled sphere of radius R, 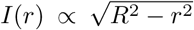, remembering that *I*(*r*) represents the z-projection of the intensity distribution. In this case, the minimum tracking error is actually found at *r* = *R*, an impractical choice. As a compromise between maximizing photon collection and minimizing tracking error, we chose *r*_*scan*_ = 3*/*4*R*.

Of course, the neuron is not actually a uniformly labeled perfect sphere; the choice of intensity distribution alters only prefactors in the gain applied to the measured center of mass location of each scan and the estimate of the measurement error. In the Kalman filter (algorithm below), these prefactors always appear in combination with quantities dependent on other parameter choices. In practice, we frequently chose *g* = 1 and adjusted other parameters to assure smooth tracking.

### Kalman Filter

With each cycle, we obtained an estimate of the offset of the neuron from the scan location and an uncertainty in this estimate. To combine this series of uncertain measurements, we used a Kalman filter whose model parameters were the position and velocity of the neuron. We treated movement in each axis independently and used a separate filter for each. Along each axis, the state variable was represented as a two dimensional vector, *x*, the coordinate of the neuron center and *v*, the velocity along that axis. The uncertainty in the estimate was represented by a covariance matrix, *P*.

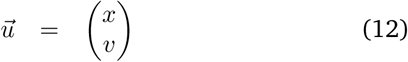

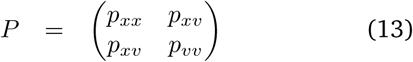

The Kalman filter consists of an update step, which propagates the model forward in time and a measurement step, which updates the model given new measurement data.

### Update step

We ran the update step with each FPGA cycle, at 40 MHz. Thus, in deriving the update rules below, we ignore terms of order Δ*t*^2^ or higher.

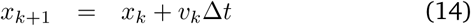

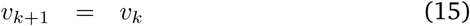

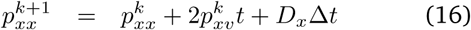

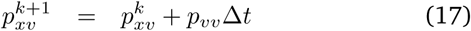

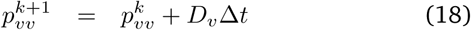

*D*_*x*_ and *D*_*v*_ represent expected diffusion in the position and velocity of the neuron; the smaller these values, the smoother the expected path and the more past measurements are factored in to the current location estimate. *D*_*x*_ and *D*_*v*_ were chosen by a combination of simulation and trial-and-error.

### Measurement step

With each complete cycle, we form an estimate of the offset of the neuron from the scan center and the variance in this estimate (see Feedback Relations above). We incorporate the new measurements independently on each axis as follows, with the superscript (*−*) indicating the pre-measurement-step values.

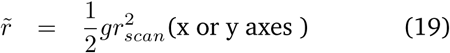

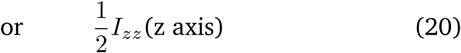

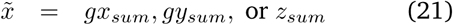

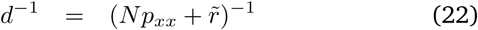

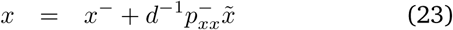

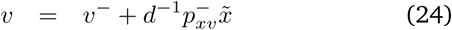

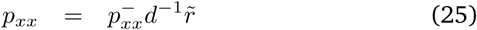

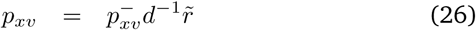

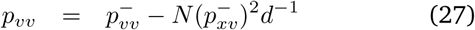

*g* and 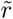 are calculated on the computer prior to tracking and passed as fixed parameters to the FPGA. Computation of *d^−^*^1^ requires multiple FPGA cycles; the entire measurement step takes about 1 μs.

The tracker can also be run with a purely spatially diffusive prior. In this case *v, D*_*v*_, *p*_*xv*_, and *p*_*vv*_ are all held fixed to 0. We often chose this mode for the z tracker, as the small range of movement imposed by the spacing between the PDMS and coverslip in the microfluidic device limited achievable axial velocities.

### Multiple Neuron Tracking

To track two neurons, we tracked the first neuron an integral number of cycles, *N*_*track*_, then switched the target of the scan to the next neuron, waited *N*_*delay*_ cycles without feedback for the mirrors to reach the new position, and then tracked the second neuron for *N*_*track*_ cycles, before switching back and repeating the cycle. For the experiments described in Figure 4 and Figure 5g, *N*_*track*_ = 4 and *N*_*delay*_ = 2, which means each neuron was tracked for 1.43 ms out of every 4.29 ms. While a neuron was not being tracked, we updated its position using the last estimated velocity, but because of limited FPGA resources, we did not update the uncertainty (P). The midpoint between the neurons was used as the input to the stage and objective piezo feedback loops.

### Calibration

To carry out the calibration experiments of Figure 1, we immobilized a larva expressing hexameric GFP and mCherry in its motor neurons between a coverslip and a slide mounted on a 3-axis piezo driven stage (MDT630B, Thorlabs). We drove the open-loop controller with a sinusoidal wave from a function generator to generate oscillations with amplitudes of 10 μm, the largest supported by the 20 μm travel of the piezo stage. Because the controller was open-loop, we did not have an independent record of the position of the stage vs. time. The amplitude and frequency of the oscillation were determined by our input to the function generator; we fit the phase to the measured path for each oscillation to determine the true location of the neuron vs. time. The tracking error (Figure 1b) represents the rms difference between the measured path and this fit.

In the axial (z-) oscillation experiments, the objective piezo scanner lagged behind the tracked position of the neuron by 8 to 13 ms. For oscillations above 10 Hz, this delay comprised a significant fraction of the oscillatory cycle; at 20 Hz, the piezo was 90 degrees out of phase and at 50 Hz, the piezo feedback was 180 degrees out of phase. In these cases, the tracking performance would have been improved by disabling the piezo feedback, but we chose to leave the feedback enabled to match experimental conditions.

### Video Recording Behavior

Larvae were recorded from below using a Basler aca640-90um CMOS camera and a 50 mm focal length c-mount lens (MVL50TM23, Thorlabs) used at a greater distance from the sensor for increased magnification. Darkfield infrared illumination was provided by the collimated output of an 850nm fiber coupled infrared led (M850F2, Thorlabs) aimed at an oblique angle from above the larva to penetrate beneath the objective. An 850 nm bandpass filter (FBH850-40, Thorlabs) was placed in front of the lens to attenuate the incoming light from the Ti:Sa laser.

### Integrating Blue Light Stimulus

Blue light stimulus was provided by the collimated output of a 450 nm laser (LP450-SF15 Thorlabs) directed from the side above the larva. We inserted a 473 nm longpass filter (LP02-473RU, Semrock) behind the objective to block transmission of this light to the photomultiplier tubes. However, we found that even with filters in place, turning on the blue light increased the green PMT count rate by an amount comparable to the recovered GCaMP6f fluorescence. This signal was likely due to autofluorescence or phosphorescence in the larval tissue, as it required the presence of a larva beneath the objective. We therefore modulated the laser to only turn on when the TAG lens was at the extremes of its focus cycle – a time when we were not using the PMT signals for either measurement or feedback. The combination of filtering and out-of-phase light presentation was sufficient to eliminate cross-talk between the blue light presentation and measurements of neural activity. Timing the stimulus light to the phase of the TAG lens meant that when the stimulus was on, the blue light flickered on and off at 140 kHz, far too fast for the larva to detect.

The approximately 5mm diameter stimulus laser beam had a power density of 0.05 W/m^2^ measured transverse to the beam. Because the light entered at an oblique angle between the objective and the larva and was subject to scattering and reflection, it is difficult to precisely estimate the power density of the light at the larva’s photoreceptors.

### Data Analysis

Videos of behavior were saved as avi files, and tracking microscope output containing position of the tracker, position of the stage, and photon counts, was written to a text file. At the beginning and end of tracking, the tracker flashed off the infrared illumination used by the video camera, allowing synchronization of the video and the tracker to within the frame rate of the camera. The Matlab function adapthisteq was used to enhance the contrast of the video images. Data were further analyzed in Matlab.

The tracker recorded the number of red and green photons collected with each revolution of the spot through a neuron. These counts had significant variation due to Poisson statistics. To estimate the underlying rates, we used a Stochastic Point Process Smoother [57] with the following model

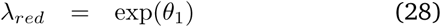

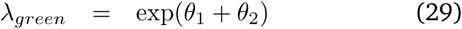

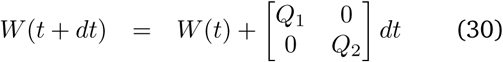

where *W* is the covariance matrix (uncertainty) for the estimates of *θ*_1_ and *θ*_2_, and *Q*_1_ and *Q*_2_ represent a prior belief in how rapidly the parameters *θ*_1_ and *θ*_2_ will change. The ratiometric measure of activity is given by exp(*θ*_2_). We chose 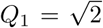 and 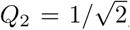, reflecting a belief that the variation in recorded intensity due to factors other than calcium dynamics should be faster than the variation due to calcium dynamics. The choice of *Q*_1_ and *Q*_2_ sets the bandwidth of the filter, but otherwise has limited impact on the results in this work. Other methods for estimating the ratio, like low-pass filtering the green and red signals then dividing, also produce similar results.

#### Cross-covariance

The normalized cross-covariance shown in Figure 4d is calculated as

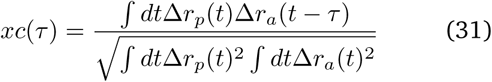

where Δ*r*_*p*_(*t*) and Δ*r*_*a*_(*t*) represent the deviation from the mean ratio for the posterior and anterior neurons respectively.

#### Ratiometric baseline correction

The red and green indicators bleached at different rates, causing a long duration shift in the ratiometric intensity baseline. In a typical recording, after 15 minutes of tracking a single neuron, mCherry fluorescence was 40% of its initial value, and GCaMP6f was not measurably bleached. To correct for this, we found the ratiometric baseline by fitting the ratiometric measure to an exponential function (*r*_*base*_ = *a* exp(*bt*)) using a truncated cost function that discards large upward deviations. The baseline corrected ratiometric measure shown in all figures (ratio/baseline) is the instantaneous estimate of the ratio divided by this baseline.

## Supplemental Methods

### Relation between locations of photon detections and displacement of the neuron

We derive an estimate of the neuron’s position and the uncertainty in this estimate for a cylindrical scan and arbitrary radial intensity distributions, extending the derivation in Berglund and Mabuchi[86]. We describe the neuron as a distribution of fluorescent label. Given fixed laser power and properties of the microscope optical train, the expected rate at which photons will be captured by the PMTs is a function of the position of the focal spot within the neuron, 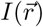.

We scan the laser beam in a cylindrical path, 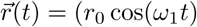, *r*_0_ sin(*ω*_1_*t*), *Z*_*A*_ cos(*ω*_2_*t*)), with *ω*_2_ ≫ *ω*_1_, *r*_0_ *< r*_*neuron*_, Z_A_ ≫ *r*_*neuron*_. The first two coordinates (*x, y*) are in the focal plane of the objective and the third *z* is aligned along the axis. For convenience *ω*_2_ is large integer multiple of *ω*_1_ (*ω*_2_ = 50*ω*_1_ in this work). As the laser beam moves through the sample, we record a sequence of photon arrivals. 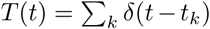 represents the rate of photon arrivals, where *t*_*k*_ is the time of arrival of the *k*^*th*^ photon. *T* (*t*) obeys Poisson statistics, so

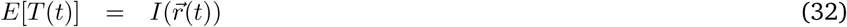

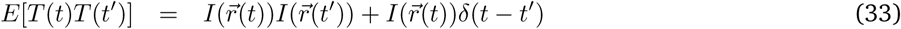

The basis for our estimation of the neuron’s displacement is a calculation at the end of each cycle of the sum of the position of the focal spot at the times when photons were emitted;

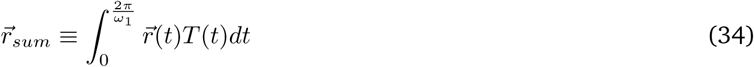

The expected value of this sum is given by

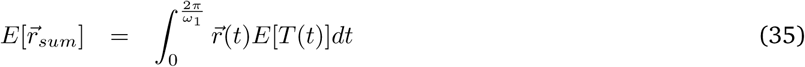

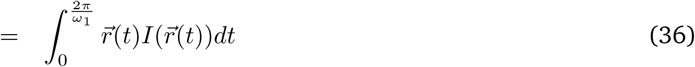

We will now calculate how this expected value depends on the displacement of the neuron from the origin of the scan. We will assume that the intensity distribution has cylindrical symmetry, that is 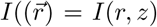. We will first consider a displacement in *x* (or equivalently *y*), then a displacement in *z*. We will consider a scan centered about the origin, with the intensity distribution of the neuron centered a distance Δ*x* away along the *x*-axis.

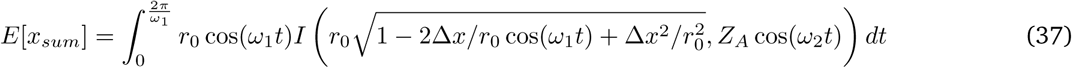

Because the axial (*z*) oscillations are much faster than the in-plane rotation of the focal spot, in the above integral, we willl replace the intensity *I*(*r, z*) with the average intensity over a *z−* scan at a given radial position

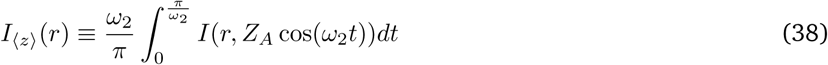

Using this approximation, and assuming a small displacement, Δ*x ≪ r*_0_, we find

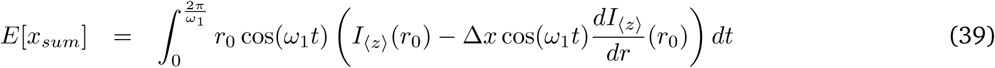

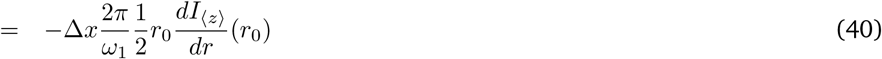

The expected number of photons collected during the scan is 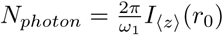. Using this relation, we can define an estimator for the *x* displacement.

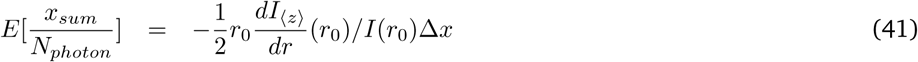

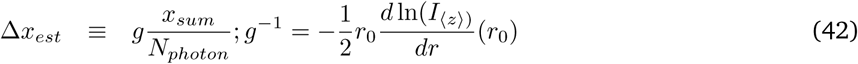

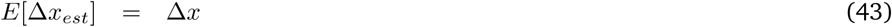

We can now calculate the variance in this estimate, 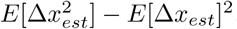. To do this, we will need to calculate the related quantity

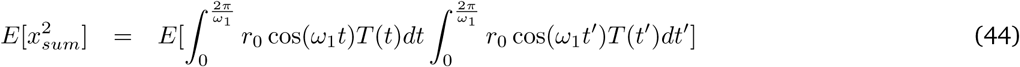

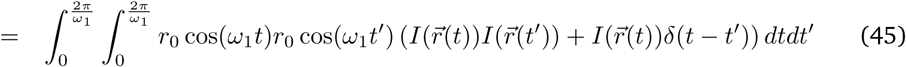

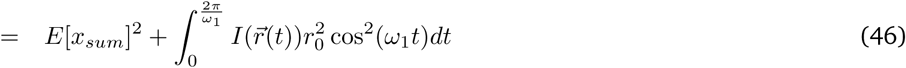

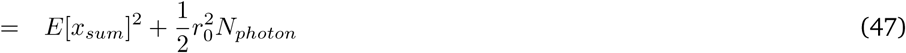

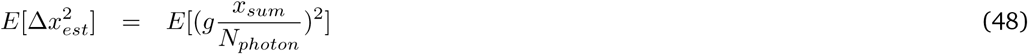

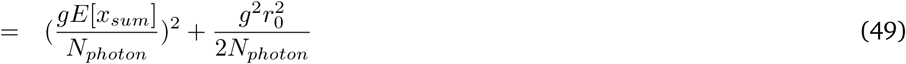

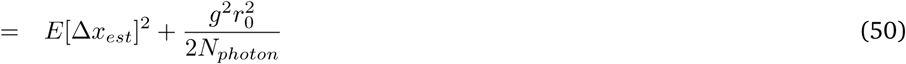

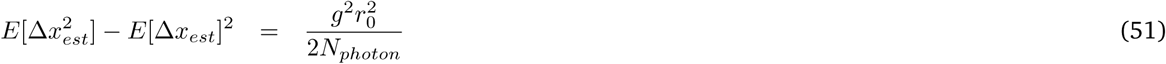

The calculation for *z* displacement proceeds along similar lines. We again use the fact that the *z*-oscillation is much faster than the rotation to write

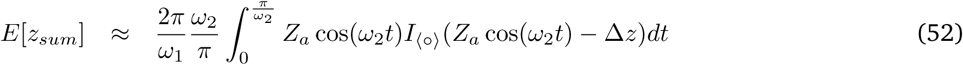

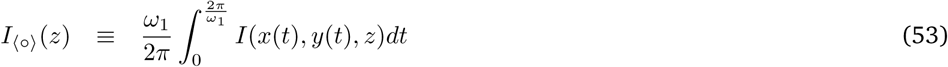

We assume that the z-scan extends well beyond to the neuron, hence the z-displacement in time is linear over the portion of the scan with appreciable fluorescence, *z ≈ Z*_*a*_*ω*_2_*t*, and we extend the limits of the integration to infinity.

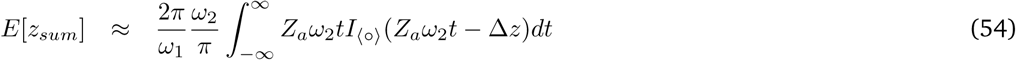

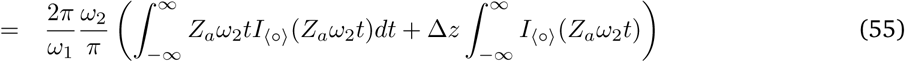

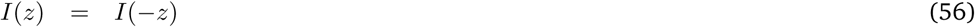

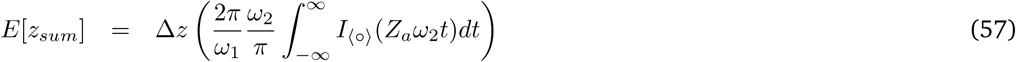

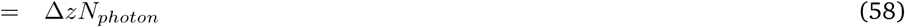

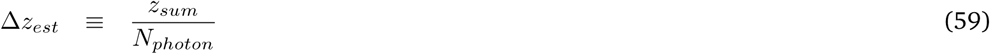

Following the same methods as for the *x* computation, we find that

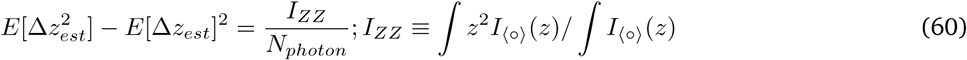

## Supplemental Figures

**Figure S1:**
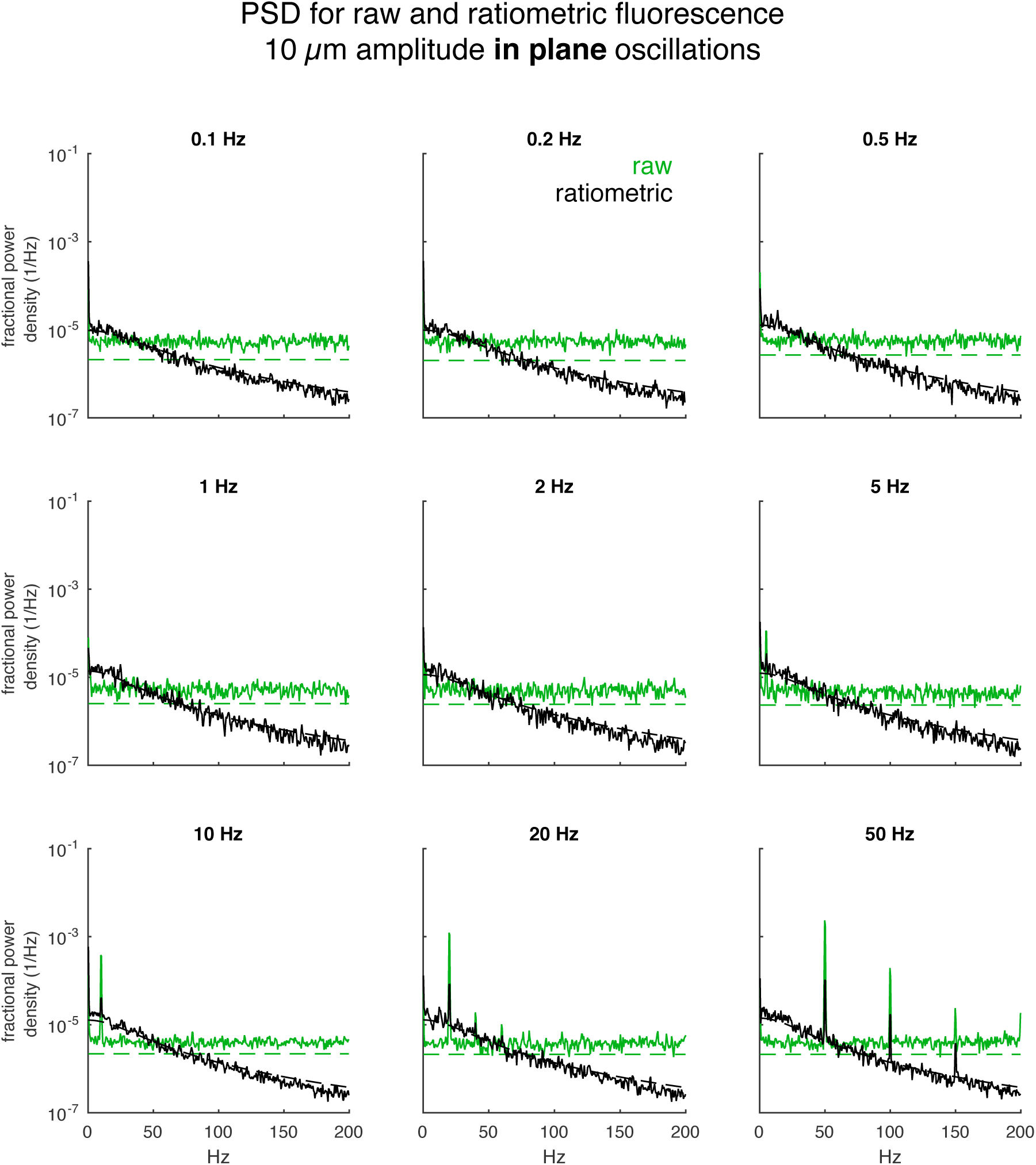
PSD for fluorescence signals from neuron sinusoidally oscillating in plane. Accompanies Figure 1. A single neuron was tracked while being oscillated in the focal plane in a sinusoidal wave with 10 μm amplitude (20 μm peak-to-peak) at varying frequencies. The green traces represent the fractional noise density for the raw green fluorescence measurement, the black traces represent the fractional noise density for the ratiometric measurement, and the dashed lines represent the expected power densities due to shot noise only. The frequency of oscillation is indicated above the graph. The maximum speed is 63 (20π) μs times the indicated frequency in Hz. The maximum acceleration is 0.39 mm/s^2^ times the square of the indicated frequency in Hz.

**Figure S2:**
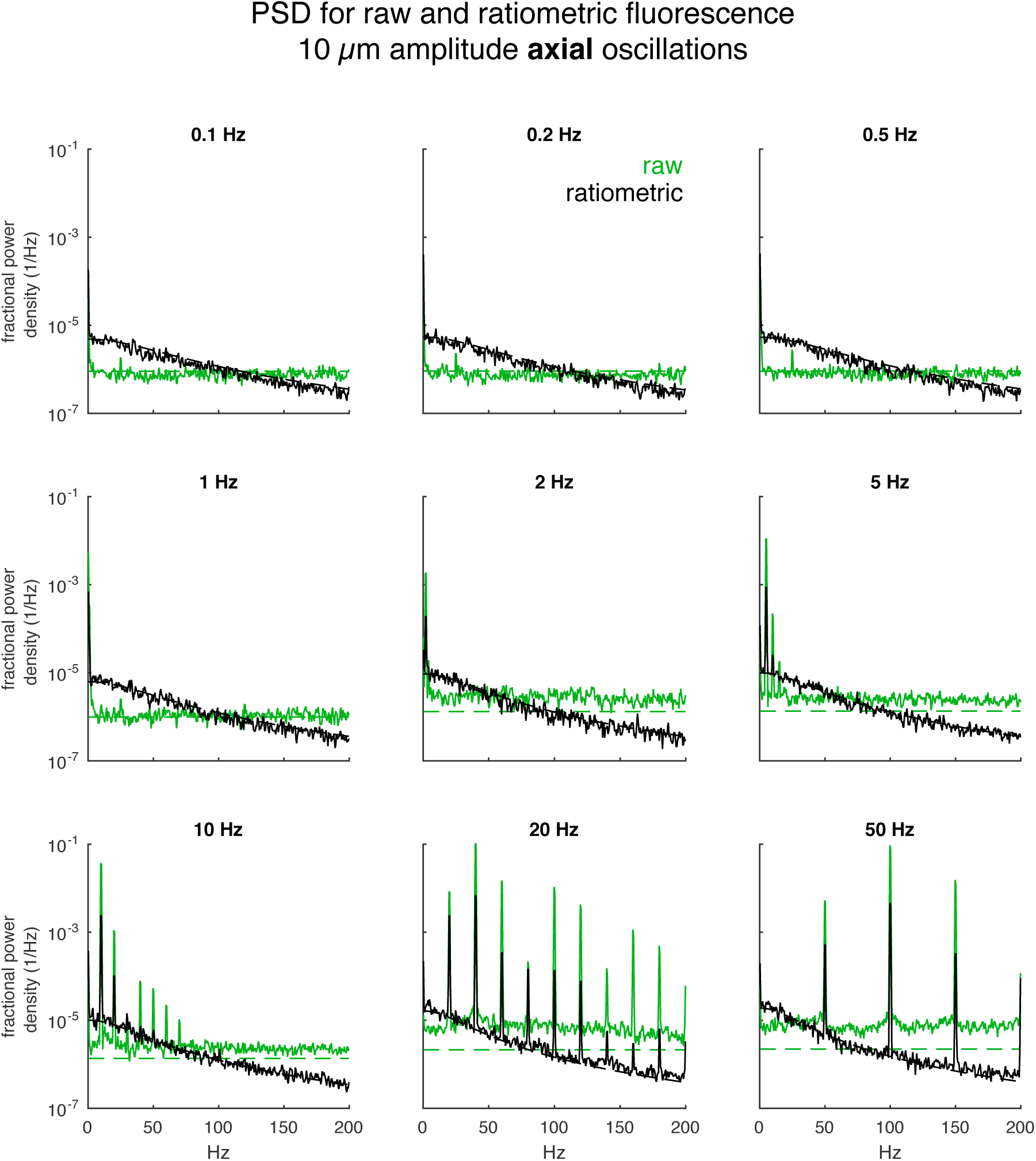
PSD for fluorescence signals from neuron sinusoidally oscillating axially. Accompanies Figure 1. A single neuron was tracked while being oscillated perpendicular to the focal plane in a sinusoidal wave with 10 μm amplitude (20 μm peak-to-peak) at varying frequencies. The green traces represent the fractional noise density for the raw green fluorescence measurement, the black traces represent the fractional noise density for the ratiometric measurement, and the dashed lines represent the expected power densities due to shot noise only. The frequency of oscillation is indicated above the graph. The maximum speed is 63 (20π) μs times the indicated frequency in Hz. The maximum acceleration is 0.39 mm/s^2^ times the square of the indicated frequency in Hz.

**Figure S3:**
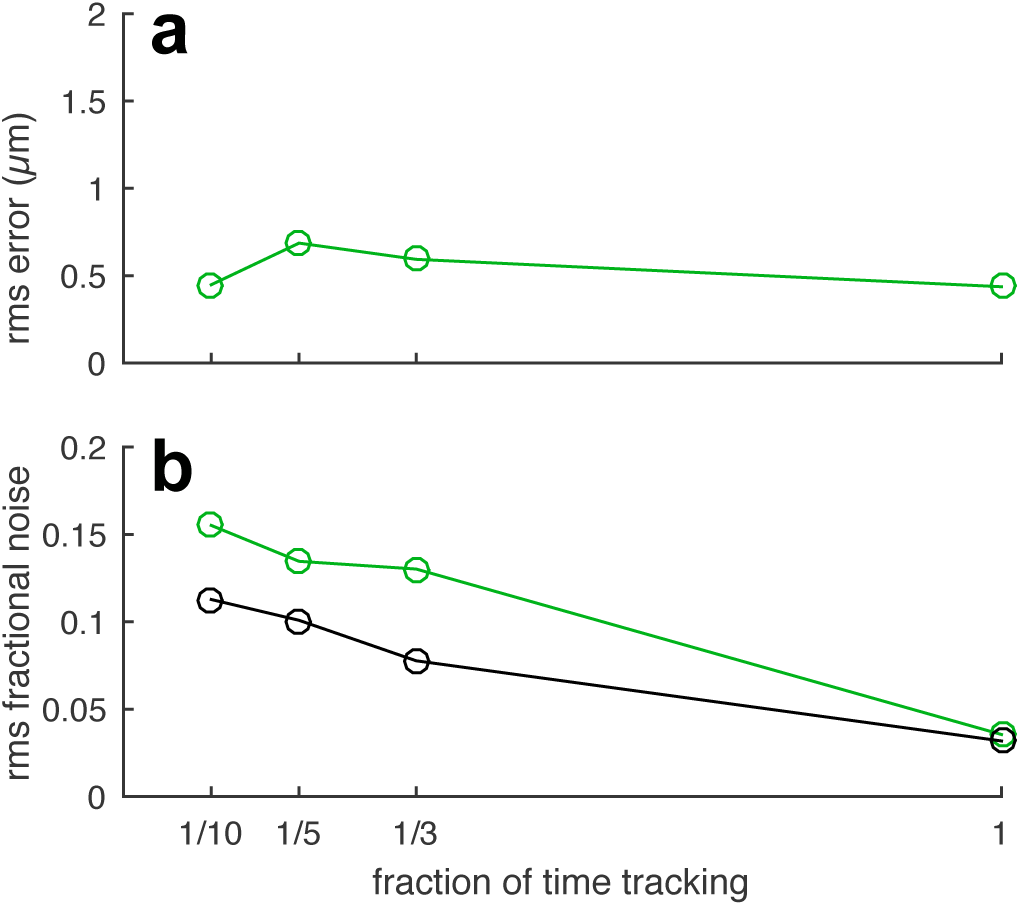
Tracking error and fluorescence noise vs. tracking duty cycle. Accompanies Figure 1. For a 10 Hz 20 μm peak-to-peak in-plane sinusoidal oscillation, the root-mean-squared (rms) positional error (**a**) and rms noise in fluorescence (**b**), if only indicated fraction of cycles are used for measurement and tracking.

**Figure S4:**
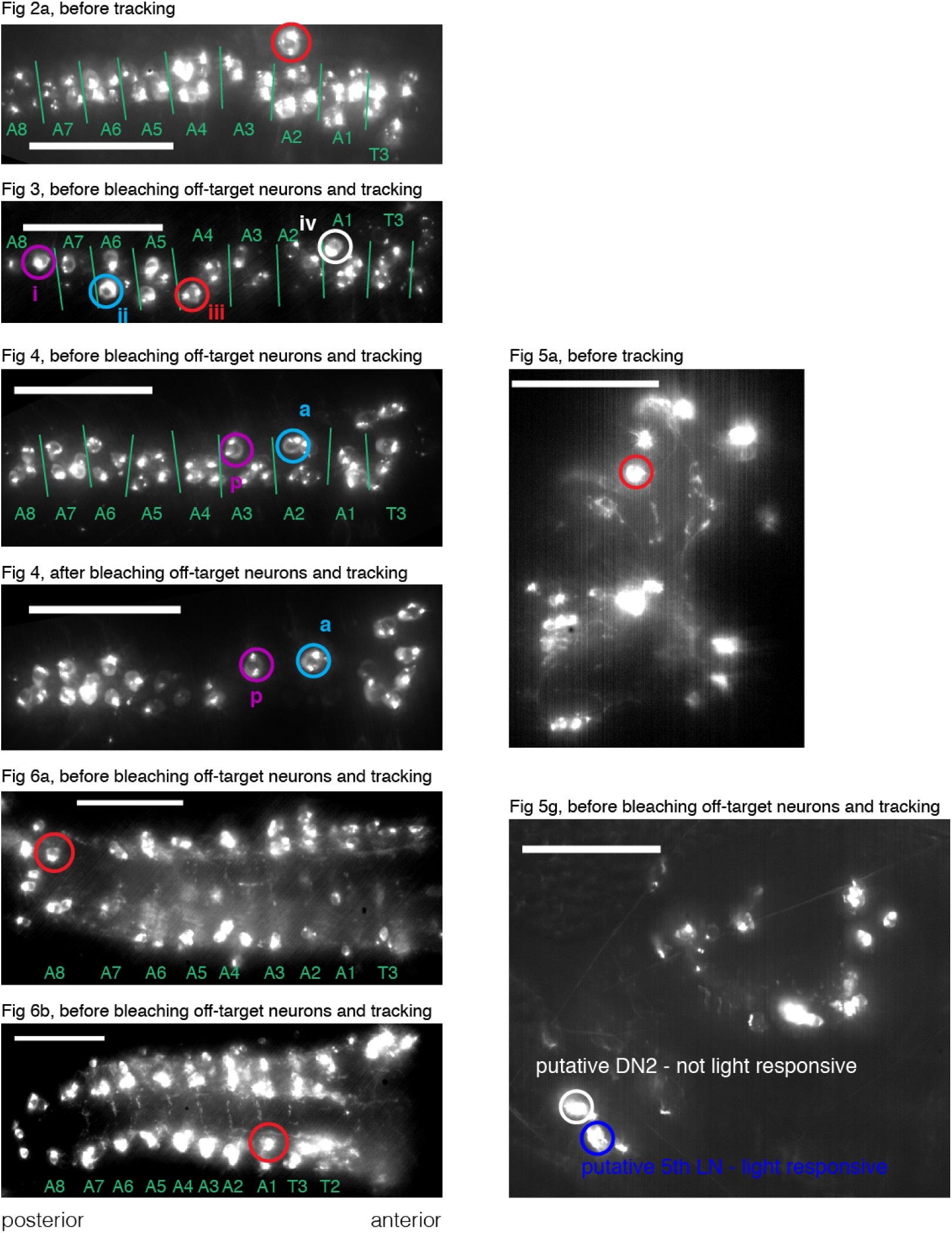
Epifluorescence images of tracked neurons. Epifluorescence images of mCherry labeling of neurons. Neurons whose activities are shown in the corresponding figures are indicated by colored circles. Images represent a maximum intensity projection of images taken at several depths, with brightness and contrast adjusted in ImageJ. Clustering of mCherry can be seen in these images; this clustering did not affect our ability to track neurons. These images were all taken while the larva was immobilized by compression, which could distort brain geometry relative to the uncompressed state. Fig 2a,3,4,6a,b – annotations mark our estimate for segment locations, based on cell body positions. Fig 6a,b: RG3602 also labels a second set of VNC neurons in a different plane from the neurons shown and tracked here. All images: scale bar is 50 μm.

## Supplemental Movies

**Supplemental Movie 1:**
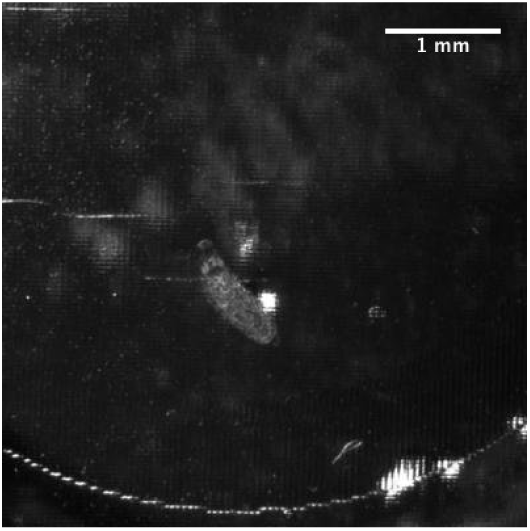
Reversible immobilization of a larva using microfluidic compression. Stereoscope recording of a larva crawling in the microfluidic device. At two points in the movie, vacuum was applied to the device, compressing and temporarily immobilizing the larva. The video was recorded at 4 frames per second; the default playback speed of 25 fps represents 6.25x real time.

**Supplemental Movie 2:**
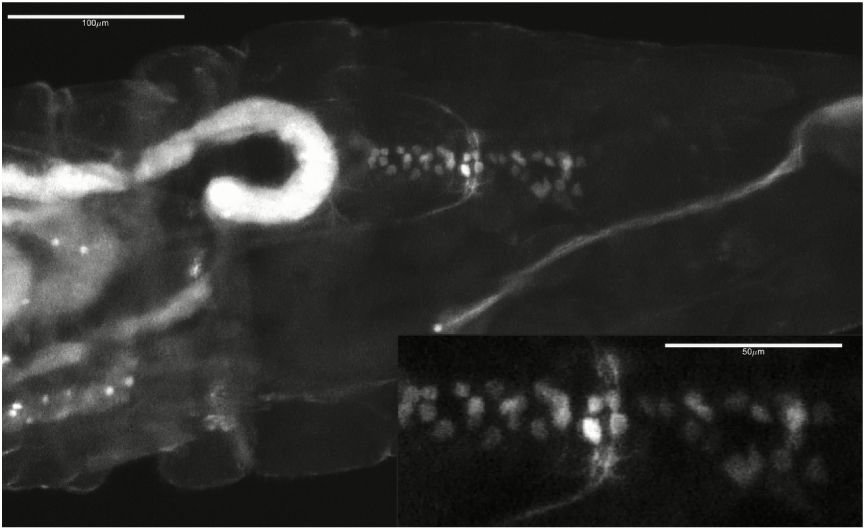
Stereoscope recording of semi-immobilized larva expressing GCaMP6f in motor neuron (RRAF > GCaMP6f, 6xmCherry). Video shows movement of a larva and time-varying fluorescence in GCaMP6f expressing VNC motor neurons as recorded through a GFP filter set. One wave of activity propagates from posterior (left) to anterior (right). In the anterior of the VNC, activity in motor neurons can be seen to correspond to contraction of corresponding segments. The inset view shows a magnified and stabilized image of only the VNC. Video was recorded at 4 frames per second; the default playback speed of 12 fps represents 3x real time.

**Supplemental Movie 3:**
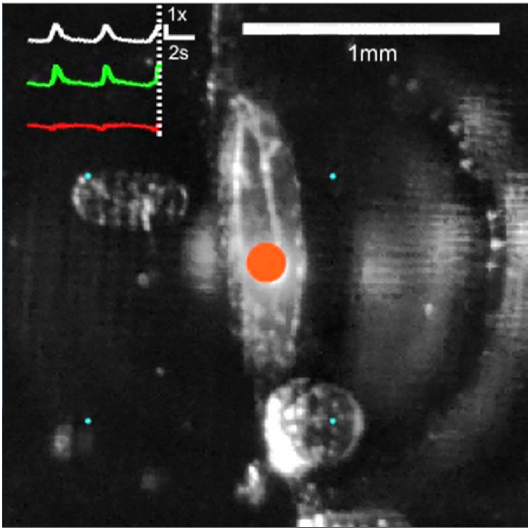
GCaMP6f in motor neuron. (RRAF > GCaMP6f, 6xmCherry; accompanies Figure 2a). This movie shows an image of the larva recorded from below (objective is out of focus above the larva). Overlaid are traces representing signals recorded from the tracked motor neuron: (from bottom to top) the red fluorescence (from hexameric mCherry, the baseline for the ratiometric measurement), the green fluorescence (from GCaMP6f, a calcium indicator), and the ratio of green to red fluorescence (ratiometric indicator of activity). The red and green traces are divided by the median signal; the ratiometric measure is divided by a baseline (low-level of activity). The infrared excitation laser beam creates an artifact visible in the camera images as a bright white spot. This spot, which does not reflect fluorescence emission from the targeted neurons or the size of the focal spot, is overlaid with a colored spot whose size and brightness indicate the instantaneous value of the ratiometric activity measure. The tracking microscope continually moves the stage to keep the targeted neuron in the center of the objective. Because of this, the crawling larva may appear stationary. To aid in the visualization of the motion, cyan dots on a 1mm grid are added to provide a fixed reference frame. Video was recorded at 10 fps; the default playback speed of 30 fps represents 3x real time.

**Supplemental Movie 4:**
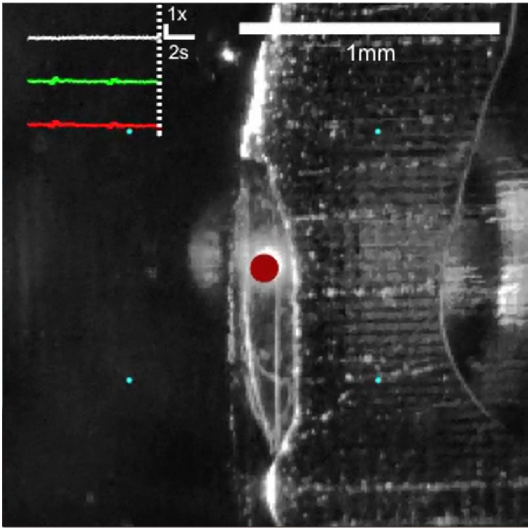
GFP in motor neuron. (RRK > 6xGFP, 6xmCherry; accompanies Figure 2f). This movie shows the behavior of a larva and measurements from a tracked motor neuron labeled with two stable proteins, hexameric GFP and hexameric mCherry. The video is annotated as for Supplemental Movie 3. The only difference is that because GFP does not respond to calcium concentration, variations in the ratiometric measure would represent motion artifacts. Note that all scales are the same as in Supplemental Movie 3. Video was recorded at 10 fps; the default playback speed of 30 fps represents 3x real time.

**Supplemental Movie 5:**
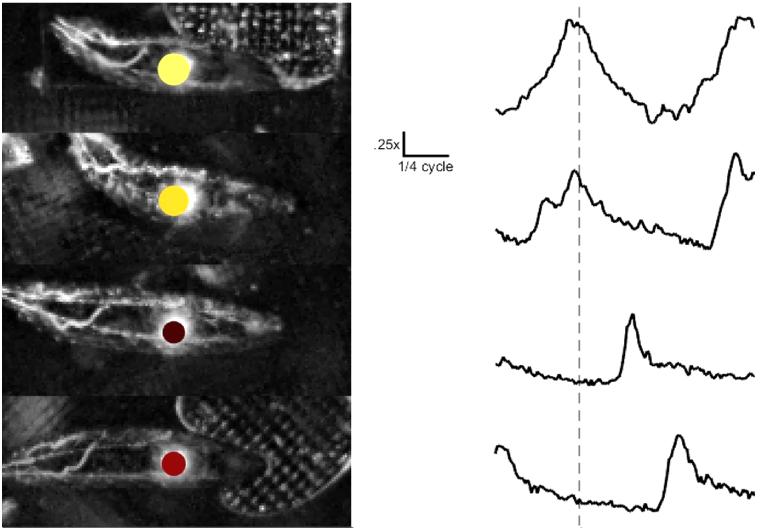
Aligned recordings from 4 motor neurons. (RRAF > GCaMP6f, 6xmCherry; accompanies Figure 3). On the left are movies of the same larva recorded at different points in time when different neurons were being tracked. The larvae have been rotated so that their paths of motion are to the right. As in Supplemental Movie 3, the size and color represent the instantaneous value of the ratiometric activity indicator. The traces to the right show the time evolution of the ratiometric indicator with the dashed line representing the current time in the video. The movies and traces have been arranged in order of neuron position in the VNC so that the most posterior neuron is at the top and the most anterior is at the bottom. The time axis for the videos and the plots has been aligned to the peristaltic cycle using measurements of neuron position from the tracker, as described in the text and Figure 3. Neither the videos themselves nor the ratiometric measures were used to produce this alignment. The video was recorded at 10 fps, and default playback is 10 fps; because the temporal axis has been warped to make the peristaltic frequency constant, playback of each larva is sometimes faster or slower than real time.

**Supplemental Movie 6:**
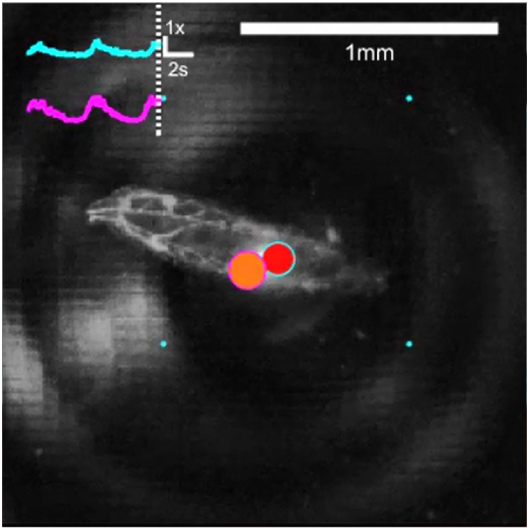
Simultaneous recording from 2 nearby motor neurons. (RRAF > GCaMP6f, 6xmCherry; accompanies Figure 4). The annotation of this video is similar to Supplemental Movie 3 except that only the ratiometric activity indicator is plotted, with the cyan line representing the more anterior of the neuron pairs. The size and color of the two spots overlaying the white laser spot represent the instantaneous activity of the neurons. The location of the spots gives the approximate orientation of the neurons relative to each other, but the displacement is greatly exaggerated (in reality both neurons would fall on the same pixel in the video image). Video was recorded at 10 fps; the default playback speed of 30 fps represents 3x real time.

**Supplemental Movie 7:**
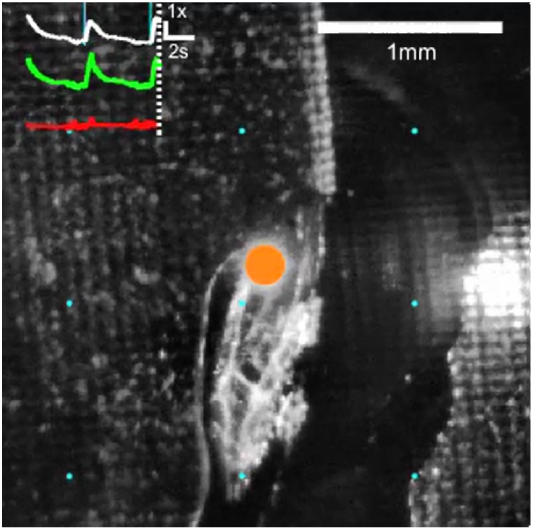
Recording from 5th LaN while presenting blue light stimulus. (tim-gal4,cry-gal80 > GCaMP6f, 6xmCherry; accompanies Figure 5a). This movie shows the behavior of a larva and activity in its 5th LaN while brief (100 ms) pulses of blue light are presented as a visual stimulus. The video is annotated as for Supplemental Movie 3, with an additional notation on the video and traces when the stimulus is on. Video was recorded at 10 fps; the default playback speed of 30 fps represents 3x real time.

**Supplemental Movie 8:**
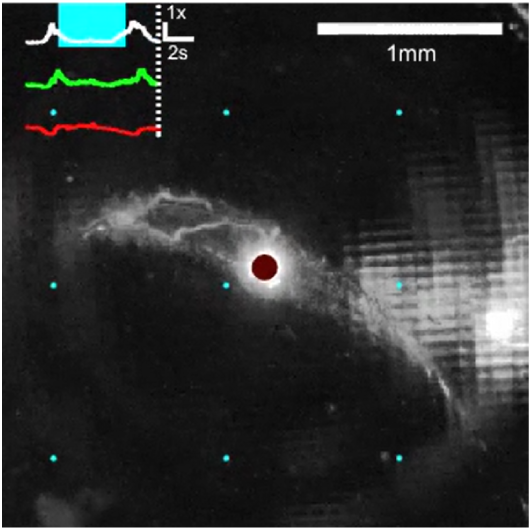
Recording from motor neuron while presenting blue light stimulus. (RRAF > GCaMP6f, 6xmCherry; accompanies Figure 5d). This movie shows the behavior of a larva and activity in an aCC or RP2 motor neuron, while longer (5 s) pulses of blue light are presented as a visual stimulus. The video is annotated as for Supplemental Movie 3, with an additional notation on the video and traces when the stimulus is on. Video was recorded at 10 fps; the default playback speed of 30 fps represents 3x real time.

**Supplemental Movie 9:**
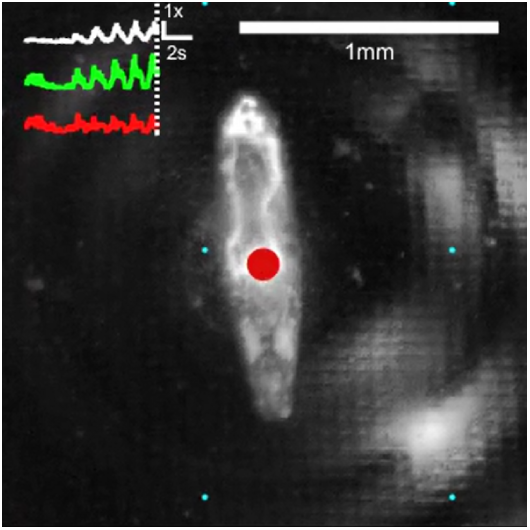
Recording from posterior A27h premotor neuron during single backward to forward transition. (R36G02 > GCaMP6f, 6xmCherry accompanies Figure 6a). This movie shows the behavior of a larva and measurements of activity in a posterior A27h premotor interneuron, for the period of time shown in the inset of Figure 6a. The video is annotated as for Supplemental Movie 3. Video was recorded at 10 fps; the default playback speed of 30 fps represents 3x real time.

**Supplemental Movie 10:**
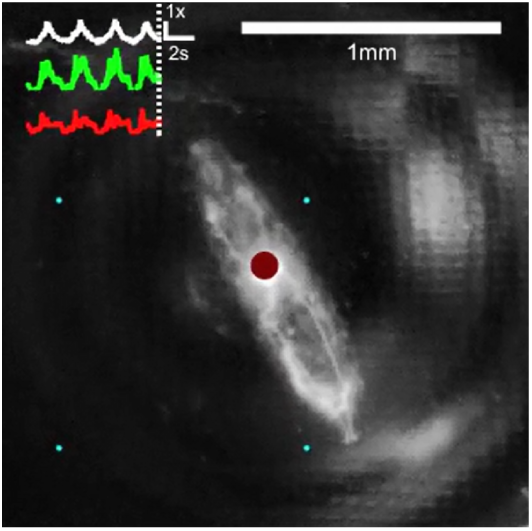
Recording from posterior A27h premotor neuron during multiple transitions. (R36G02 > GCaMP6f, 6xmCherry accompanies Figure 6a). This movie shows the behavior of a larva and measurements of activity in a posterior A27h premotor interneuron, for the full period of time shown in Figure 6a. The video is annotated as for Supplemental Movie 3. Video was recorded at 10 fps; the default playback speed of 30 fps represents 3x real time.

**Supplemental Movie 11:**
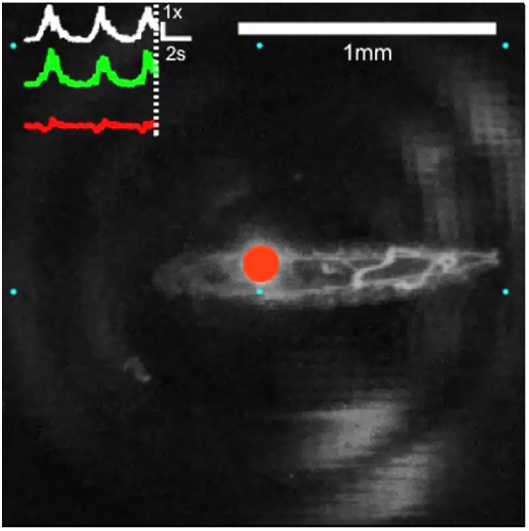
Recording from anterior A27h premotor neuron during single forward to backward transition. (R36G02 > GCaMP6f, 6xmCherry accompanies Figure 6b). This movie shows the behavior of a larva and measurements of activity in an anterior A27h premotor interneuron, for the period of time shown in the inset of Figure 6b. The video is annotated as for Supplemental Movie 3. Video was recorded at 10 fps; the default playback speed of 30 fps represents 3x real time.

**Supplemental Movie 12:**
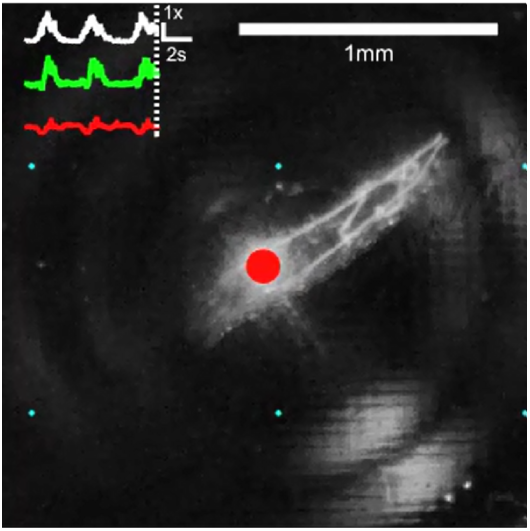
Recording from anterior A27h premotor neuron during multiple transitions. (R36G02 > GCaMP6f, 6xmCherry accompanies Figure 6b). This movie shows the behavior of a larva and measurements of activity in an anterior A27h premotor interneuron, for the full period of time shown in Figure 6b. The video is annotated as for Supplemental Movie 3. Video was recorded at 10 fps; the default playback speed of 30 fps represents 3x real time.

